# Chemically Induced Senescence in Human Stem Cell-Derived Neurons Promotes Phenotypic Presentation of Neurodegeneration

**DOI:** 10.1101/2021.07.11.451956

**Authors:** Ali Fathi, Sakthikumar Mathivanan, Linghai Kong, Andrew J Petersen, Cole R. K. Harder, Jasper Block, Julia Marie Miller, Anita Bhattacharyya, Daifeng Wang, Su-Chun Zhang

## Abstract

Modeling age-related neurodegenerative disorders with human stem cells is difficult due to the embryonic nature of stem cell derived neurons. We developed a chemical cocktail to induce senescence of iPSC-derived neurons to address this challenge. We first screened small molecules that induce embryonic fibroblasts to exhibit features characteristic of aged fibroblasts. We then optimized a cocktail of small molecules that induced senescence in fibroblasts and cortical neurons without causing DNA damage. The utility of the “senescence cocktail” was validated in motor neurons derived from ALS patient iPSCs which exhibited protein aggregation and axonal degeneration substantially earlier than those without cocktail treatment. Our “senescence cocktail” will likely enhance the manifestation of disease-related phenotypes in neurons derived from iPSCs, enabling the generation of reliable drug discovery platforms.

## Introduction

As global life expectancy increases, neurodegenerative disorders are predicted to cause a staggering burden to society. Substantial efforts have been made to develop effective therapies, but progress is slow and drugs developed based on animal studies have so far mostly failed in clinical trials ^1, 2^. Poor clinical translatability of animal models necessitates additional models to test therapeutic strategies.

Human pluripotent stem cell (hPSC) derived neurons model early stages of neurodegeneration and have potential benefits in drug discovery and testing. An advantage of the hPSC model is the ability to capture the human genetic background underlying diseases by establishing patient specific iPSCs, or by studying specific effects of disease proteins via introducing disease-related mutations into otherwise normal hPSCs ^3, 4^. However, the iPSC reprogramming process erases many of the aging marks found in somatic donor cells ^5, 6, 7, 8^ and hPSC-derived neurons are similar to those in fetal development, based on transcriptional and functional profiling ^9, 10, 11^. Thus, generating hPSC-derived neurons that resemble those in the adult and aging brain is critical for modeling neurodegenerative diseases using hPSCs.

One approach for modeling cellular senescence is trans-differentiation of fibroblasts or other aged somatic cells into neurons, avoiding the pluripotent stage and maintaining senescence markers ^12, 13^. Indeed, a recent study showed that transdifferentiated neurons retain age related transcription profiles and manipulation of RANBP17 was able to reverse some of the age-related transcriptional changes in iPSC-derived neurons ^13^. However, direct conversion of fibroblasts into neurons is relatively low throughput given the lack of expansion capacity of the resulting neurons.

Modulation of genes linked to premature aging disorders is another strategy to accelerate aging in stem cell models. The ectopic expression of progerin, a mutant form of nuclear lamina protein A (LMNA) that causes accelerated aging in progeria, in an iPSC model of Parkinson’s disease can trigger age-related and degenerative phenotypes, including neuromelanin accumulation, dendrite degeneration, loss of tyrosine hydroxylase and accumulation of pathological aggregates ^7^. It remains to be determined how closely these approaches model physiological aging in normal neurons or pathological aging seen in late-onset diseases. Overexpression of premature aging genes introduces the challenge of distinguishing phenotypes related to the disease from those induced by foreign gene overexpression.

In the current study, we screened for chemicals/pathways that selectively trigger senescence phenotypes in primary neonatal fibroblasts and iPSC derived cortical neurons. To identify pathways important in neuronal senescence, we first used transdifferentiated neurons from aged and young fibroblasts and identified molecular markers for neuronal aging, including decreased expression of H3K9Me3, chromatin associated protein HP1γ and lamina associated polypeptide Lap2β. We then used these readouts for screening small molecules and developed a combination of molecules that induce senescence and protein aggregation in cortical neurons differentiated from hPSCs. We evaluated this chemical-induced senescence (CIS) approach in motor neurons derived from ALS (TARDBP mutant) patient iPSCs and confirmed that CIS promoted earlier and consistent manifestation of disease related phenotypes. Furthermore, using autophagy activator molecules, we were able to mitigate cellular senescence (CS) phenotypes in the MNs. Thus, this CIS strategy enables more effective iPSC modeling of phenotypes in ALS.

## Results

### Identification of small molecules that induce senescence in neonatal fibroblasts

Primary human fibroblasts retain age-related markers depending on the age of the individual from which the cells are isolated 14. These cells are thus appropriate reference for studying cellular senescence (CS). We compared neonatal fibroblasts with those from a 72-year old male and 62-year old female donors by examining the expression of age-related markers H3K9Me3, Lap2β, and HP1γ. We found that neonatal fibroblasts expressed a higher level of H3K9Me3, Lap2β, and HP1γ than old fibroblasts (72 years) in our high content imaging platform. In addition, the old fibroblasts expressed the senescence associated β-Gal (Figure 1A, 1B, S1A-C). These findings are consistent with a previous observation 7, indicating that these markers are reliable readouts for assessing CS.

**Figure 1.**
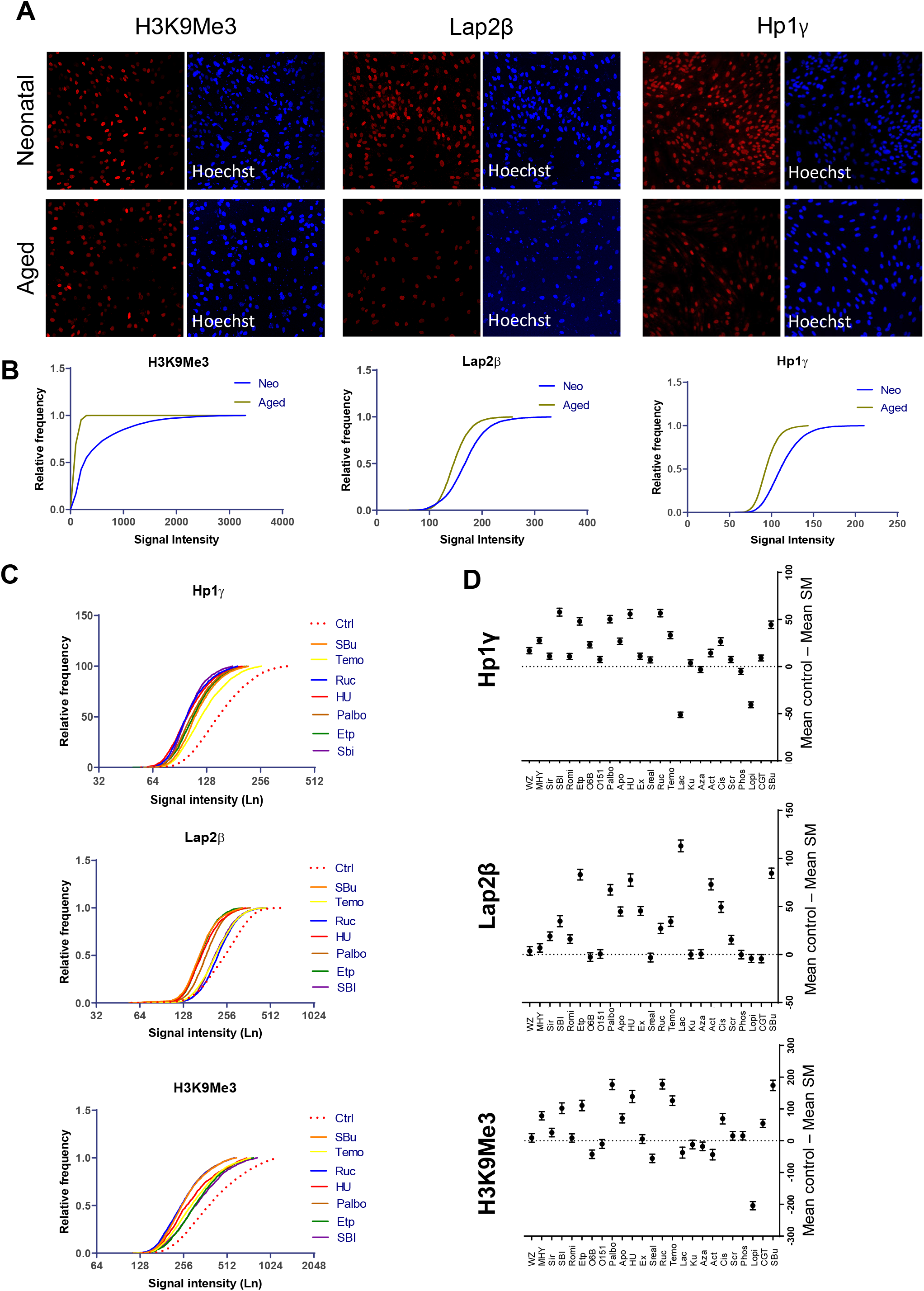
Identifying small molecules for inducing CS in human neonatal fibroblasts. Immunostaining for H3k9Me3, Lap2β and HP1γ proteins in both neonatal and aged fibroblasts Scale bar= 100μm (A). Frequency distribution analysis for different bins of signal intensity in high content imaging for H3k9Me3, Lap2β and HP1γ proteins in male neonatal and aged (72 years old) fibroblasts (B). Frequency distribution analysis for H3k9Me3, Lap2β and HP1γ protein expression in male neonatal fibroblasts treated with different small molecules; dashed red line is control and top seven molecules for each protein showed in the graph (C). Mean difference for signal intensity of all 25 small molecules depicted as mean ± 95% confidence intervals compared to the DMSO control group. The zero line means no difference compare to control and if difference in the mean does not touch the reference line then changes in expression are significant (D).

We then looked for molecules that may induce senescence phenotypes in the neonatal fibroblasts, focusing on the known senescence associated pathways. We selected 25 small molecules known to affect pathways involved in CS ^15^, including autophagy related molecules, Akt signaling, and inhibitors of mTOR, HDAC, ZMPSTE24, and Sirtuin signaling (Table S1). We examined the toxicity of these molecules in their minimum effective concentrations based on previous studies using calcein AM and ethidium homodimer (EthD-1) fluorescent dyes to distinguish live versus dead cells. None of the small molecules induced cell death beyond the DMSO control (5-10% cell death) at the final selected concentration (Figure S1D). By culturing the neonatal fibroblasts in the presence of the small molecules at an effective dose for 5 consecutive days and examining the expression of the above CS markers, we found that more than half of the molecules (13 molecules, p≤ 0.001, Table S2) significantly decreased the expression of all three readouts (Figure 1C, 1D). Among the 13 molecules, seven also induced expression of β-Gal, another consensus marker for CS (Figure S1E, S1F). Thus, we identified a set of small molecules that induce senescence phenotypes in neonatal fibroblasts.

### Identifying small molecule cocktails that enhance neuronal senescence

Epigenetic marks, including those associated with aging, are largely erased during reprogramming to iPSCs ^6, 16^. Consequently, cells differentiated from iPSCs, including neurons, behave like those in embryonic development. In contrast, neurons directly converted from fibroblasts by forced expression of transcription factors retain much of the age-related signatures in their parental somatic cells ^13^. To validate this phenomenon and to establish CS readouts in neurons, we reprogrammed both young and old human fibroblasts to neurons using a combination of gene overexpression and small molecules ^13^. Both neonatal and aged fibroblasts were transduced with lentiviral particles for Eto and XTP-Ngn2:2A:Ascl1 (N2A) and expanded in the presence of G418 and puromycin for at least three passages. Induced neurons (iNs), exhibiting polarized morphology and expressing neuronal proteins like β-III tubulin (Figure 2A), appeared at the 2^nd^ week in the neonatal fibroblast group and mostly at the 3^rd^ week for the old fibroblast group. At the end of 3 weeks of DOX treatment, the mean conversion rate for neonatal iNs was 18.1%±3.5, whereas for aged iNs was 39%±4.4 (Figure 2B). Importantly, the iNs from old fibroblasts showed a lower intensity in the epigenetic mark H3K9Me3, Lamin B2, and Lap2β as well as the heterochromatin protein HP1γ (Figure 2C). Besides the above markers, the morphology and size of a cell and nucleus may serve as a sign of CS^17^. We also noticed that neonatal iNs had a lower Hoechst (nuclear) intensity (Figure 2D) and a smaller nucleus area compared to their aged counterparts (Figure 2E), while there were no differences in the nuclear roundness and ratio between young iNs and aged iNs (Figure 2F, 2G). Our results confirmed that the iNs from aged fibroblasts retain the age-related signatures of their parental cells, setting a reference for us to examine the effects of small molecules on CS in embryonic neurons.

**Figure 2.**
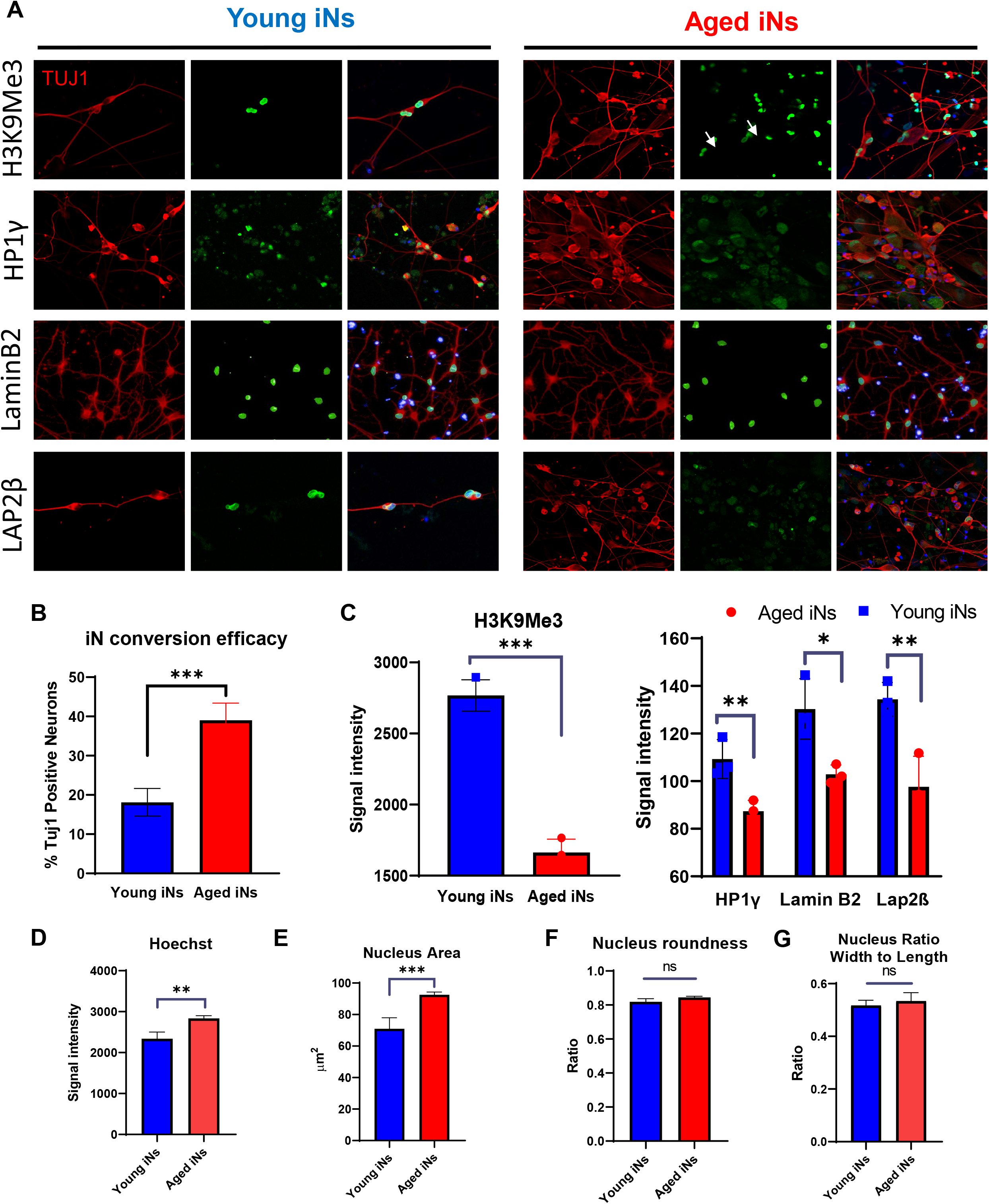
Cellular senescence marks are preserved during direct reprograming of fibroblasts to neurons. Immunostaining for H3K9Me3, LaminB2, Lap2β and HP1γ co-stained with TUJ1 (red) in induced neurons (iNs) derived from fibroblasts from both neonatal and a 72 year-age donor Scale bar= 100μm (A). Quantification results for percentage of TUJ1 positive neurons (B) and mean signal intensity for H3K9Me3, Lap2β, LaminB2 and HP1γ (C). Quantification results for Hoechst signal intensity (D), nucleus roundness (E), nucleus ratio (F) and nucleus area (G) for both young and aged iNs (ns: not significant, *: p<0.05, **: p<0.01, ***: p<0.001 unpaired t-test).

Neurons differentiated from ESCs and iPSCs resemble those during embryonic development. To identify small molecules that induce CS in the embryonic neurons, we generated cerebral cortical neurons from GFP-expressing hESCs (H9, WA09) according to an established protocol ^18^ (Figure S2). The ESC-derived cortical progenitors at day 14 expressed SOX1 (86.7%) and OTX2 (87%), markers of cortical progenitors (Figure S2B). When differentiated to mature neurons in the presence of compound E that inhibits notch signaling and MEK inhibitor PD0325901 at day 21, the majority of the cells expressed neuronal markers (MAP-2b 95%, TUBB3 95%) (Figure S2C). Following treatment of the neuronal cultures with small molecules for 4 consecutive days, we assayed for CS hallmarks (Figure S2D). The criteria for positive molecules were defined by expression of CS markers without inducing obvious DNA damage and cell death. By using three different concentrations based on the half maximal inhibitory concentrations (IC50s) for each small molecule, we identified a concentration that did not cause cell death (Figure S2E). Romidepsin, O151, SBI-0206965, Lopinavir, Sodium Butyrate, SCR-7 and Phosphoramidon had a significant impact on the expression of all three readouts H3K9Me3 (Mean±SEM 1980±22, 1957±19, 1632±15, 1806±27, 1990±18, 1908±23, 2037±24, respectively, compared to 2183±14 in control), Lap2β (742±6.4, 688±6, 726±5, 709±8, 734±5, 693±7.7, 855±7.5, respectively, compared to 789±4 in control) and HP1γ (122±3.6, 98±0.5, 92± 0.3, 96±0.64, 98±0.5, 99±0.5, 95±0.5, respectively, compared to 108±0.5 in control) (Figure 3A, 3B, Table S2). Romidepsin induced a greater expression of HP1γ and Phosphoramidon induced greater Lap2β expression compared to the mean expression in the control group and were excluded from further experiments (Table S2). Among the remaining molecules, we found that neurons treated with Actinomycin D, Etoposide, Temozolomide and Hydroxy-urea showed higher H2A.x expression compared to the control group (Figure 3C, 3D), suggesting that these molecules caused significant DNA damage, promoting us to exclude these molecules from further screenings. Five molecules were selected for further analysis (O151, SBI-0206965, Lopinavir, Sodium Butyrate, SCR-7).

**Figure 3.**
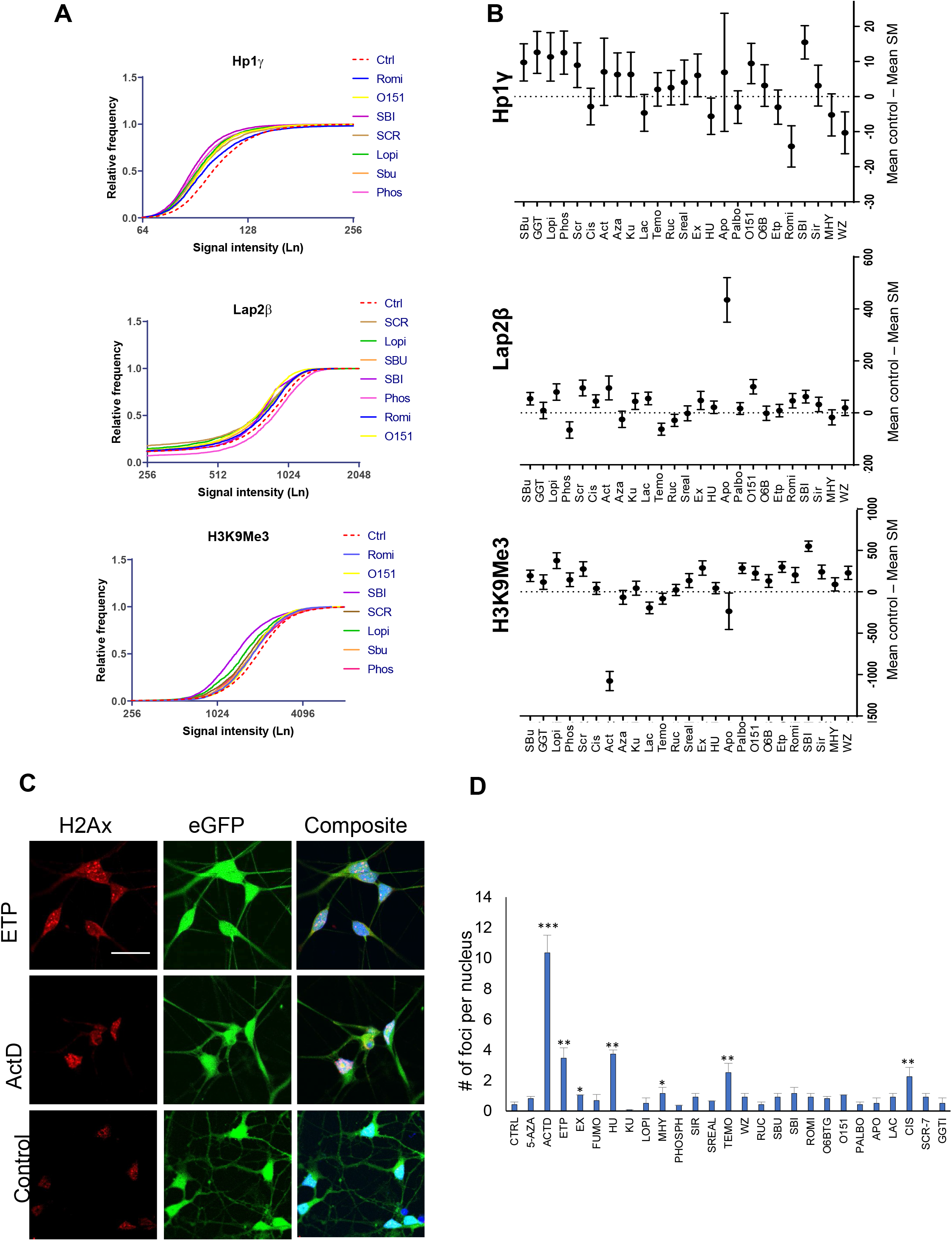
Chemical induction of CS in hESC-derived cortical neurons. Frequency distribution analysis of high content imaging data for H3K9Me3, Lap2β and HP1γ proteins in cortical neurons. The dashed red line is control and top seven molecules for each protein marker are shown in the graph (A). Mean difference for signal intensity of all 25 small molecules depicted as mean ± 95% confidence intervals compared to the DMSO control. The zero line means no difference compare to the control and if difference in the mean does not touch the reference line then changes in expression are significant (B). Confocal images of phospho-Histone H2A.X (Serine 139) in the H9-GFP cortical neurons treated with Etoposide, Actinomycin D and DMSO as control (Scale bar=50 μm) (C). Quantification results for the number of positive foci for phospho-Histone H2A.X (Serine 139) per nucleus in cortical neurons treated with different small molecules (D).

Our next step was to identify whether any combination of these five small molecules induces CS in neurons. We used the single molecule treatment with SBI-0206965 (autophagy inhibitor) as a reference since it had greater performance in modulating all three readouts during the initial screening. In this set of experiments, we used 50% of the concentration that we used for first set of experiments for molecules used in pairs and 70% reduction in triple combination to minimize cell toxicity. Results showed that most of the combinations had greater or similar effect to SBI (Figure 4A). Two of the combinations, SLO (SBI-0206965, Lopinavir, O151) and SSO (SBI-0206965, Sodium Butyrate, O151), had a greater mean difference in H3K9Me3 and Lap2β expression compared with both DMSO (Control) and SBI-0206965 treated cells (p < 0.01).

**Figure 4.**
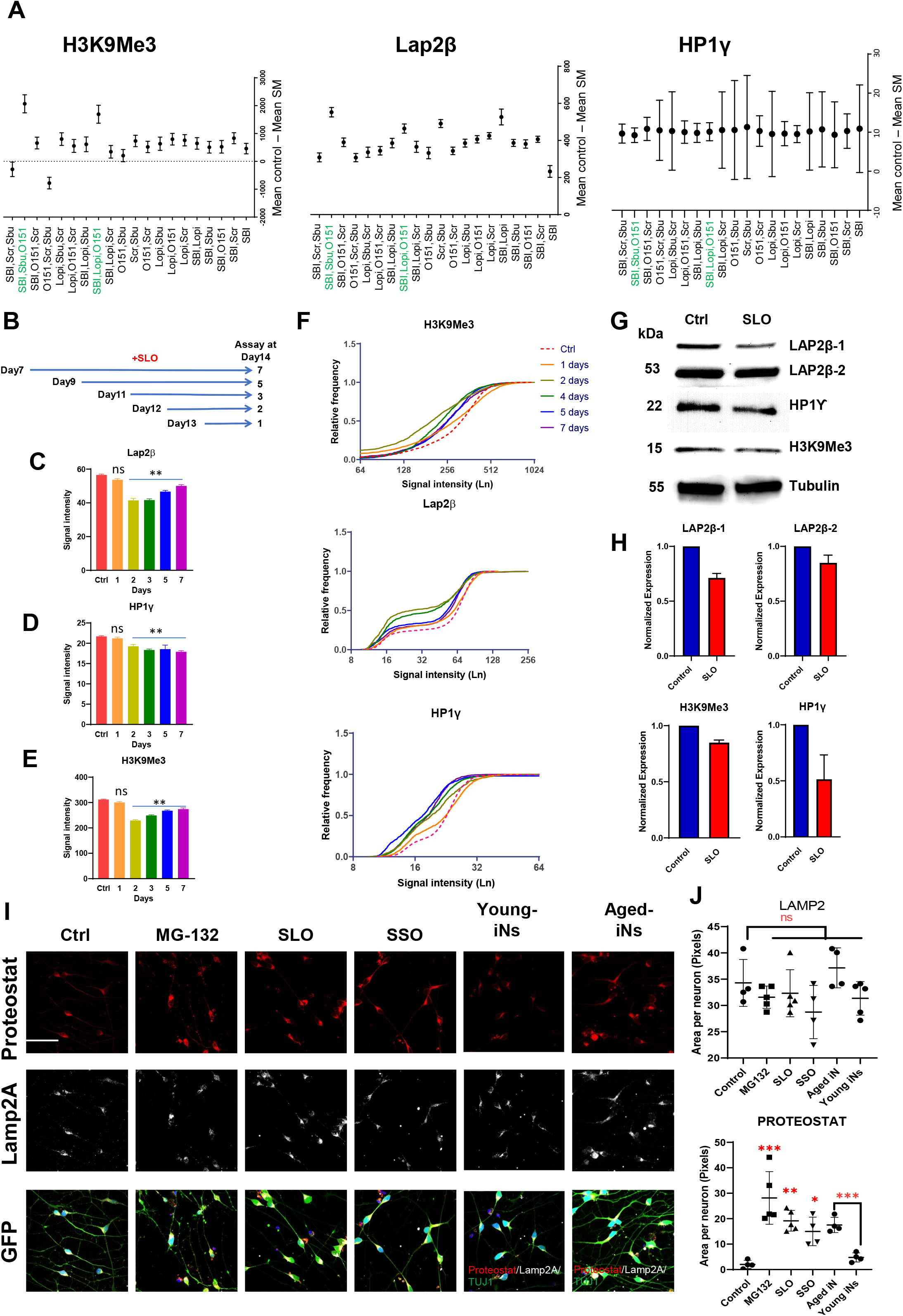
Combinatorial effect of small molecules on CS in cortical neurons. Different combination of five most effective molecules (O151, SBI-0206965, Lopinavir, Sodium Butyrate, SCR-7) tested on cortical neurons and mean expression of H3K9Me3, Lap2β and HP1γ in treatment groups compared to the DMSO control (A). A graph showing the period of SLO treatment on the expression of Lap2β, HP1γ and H3K9Me3 at day 14 after maturation (B) and high content imaging quantification of signal intensity for each marker after SLO treatment (C-E). Relative frequency distribution of different bins of signal intensity for Lap2β, HP1γ and H3K9Me3 in cortical neurons treated with different small molecules (F). Representative images of Western blot for all three markers in cortical neurons treated at day 21 of differentiation (G) and their normalized protein expression to tubulin expression (H). Immunostaining images of H9-GFP cortical neurons treated with MG-132 (proteasome inhibitor), SLO (SBI-0206965, Lopinavir and O151) and SSO (SBI-0206965, Sodium Butyrate and O151) and stained for Lamp2A (Lysosome membrane associated protein) and Proteostat dye for detection of protein aggregation (Scale bar=100 μm) (I), and quantification of positive area in neurons for Lamp2A and Proteostat (J). Young and aged iNs added for comparison with ESC-derived cortical neurons (I, J). (ns: not significant, *: p<0.05, **: p<0.01, ***: p<0.001 one-way ANOVA with Dunnett’s multiple comparison test).

To determine the minimum period of treatment needed to induce stable CS, differentiated cortical neurons at day 7 were treated with the SLO small molecules for different periods of time (treated at day 7, day 9, day 10, day 12 and day 13) and the cells were analyzed at day 14. Expression of H3K9Me3, HP1γ and Lap2β indicated that 2-4 days of continuous treatment with SLO molecules resulted in the maximum effect (Figure 4B-F). This experiment showed that expression of H3K9Me3 and Lap2β at 5- and 7-days post treatment recovered slightly but not to the normal condition. Reduction in HP1γ level was more persistent following SLO treatment and stayed at a lower level compared to the control cells even at 5- and 7-days post treatment (Figure 4B-F). Down regulation of all three senescence related proteins, H3K9Me3, HP1γ and Lap2β, was confirmed by western blot in the differentiated cortical neurons treated with SLO at day 7 (Figure 4G, H).

In addition to the CS phenotypes analyzed above, neuronal senescence is often accompanied by intracellular protein aggregation. We hence examined the effect of the top two small molecule combinations on protein aggregation with MG-132-treated cells (a proteasome inhibitor) as a positive control. Proteostat^TM^ staining revealed protein aggregates in cells treated with SSO or SLO comparable to MG-132 condition which were colocalized by Lamp2 positive autophagosomes (Tukey’s multiple comparison MG-132 p<0.0001, SLO p<0.004, SSO p<0.035) (Figure 4I, J). As additional controls, Proteostat^TM^ and Lamp2 staining revealed more prominent protein aggregation in aged iNs than in the young iNs (p<0.003). Our results show that the CS phenotype in neurons induced by small molecules is associated with intracellular protein aggregation, similar to the phenomena preserved in aged iNs.

Mitochondrial defects are associated with senescence in the directly reprogrammed neurons^19^. We found that SLO treated neurons showed a higher ROS level than the control cells, revealed by MitoSoX staining (Fig. S3A, C). It is, however, much lower than that in cells treated with FCCP, a potent uncoupler of mitochondrial oxidative phosphorylation and inducer of cell apoptosis. In parallel, JC-10 assay showed that the SLO treated cortical neurons had a lower mitochondrial membrane potential than untreated controls, but again not as low as that in the FCCP treated cells (Fig. S3B, D). Accompanying with the functional changes was morphological alterations in the mitochondria when neurons treated with SLO, including lower branches and smaller area for SLO treated cells, though statistically insignificant (Fig. S3E-G). Thus, SLO-induced CS is accompanied by functional alterations in mitochondria, including depolarization and over production of ROS.

### SLO-treated neurons express CS-related transcripts and pathways

To define CS-related changes in SLO-treated neurons, we performed RNA-seq analysis on cortical neurons treated with or without SLO. Principal component analysis based on overall gene expression showed high similarity (clustering) among independently cultured neurons treated with SLO or among those without SLO treatment (controls), but a high degree of separation between the SLO-treated and the control groups (Figure 5A). When comparing our RNAseq data with iNs from both young (<30 yrs, 8 samples) and aged (>60 yrs, 9 samples)13, we found that our cortical neurons are similar to young iNs, whereas the SLO treated neurons clustered with aged iNs (Figure 5A). We further compared our SLO treated cells to the aged (>60 yr, N=205) and young (<30yr, N=128) brain samples available in PsychENCODE20. The SLO treated samples (orange dots) are clustered with the PsychENCODE aged group (red dots), whereas the CTRL samples (blue dots) are clustered with the PsychENCODE young group (green dots). Note that we used PC2 and PC3 for showing clustering since the first PC (PC1) likely captures potential major confounding factors between our study and PsychENCODE (Figure 5B). Together, our results indicate that the SLO-treated neurons resemble those in the aged human brain and those directly converted from aged fibroblasts.

**Figure 5.**
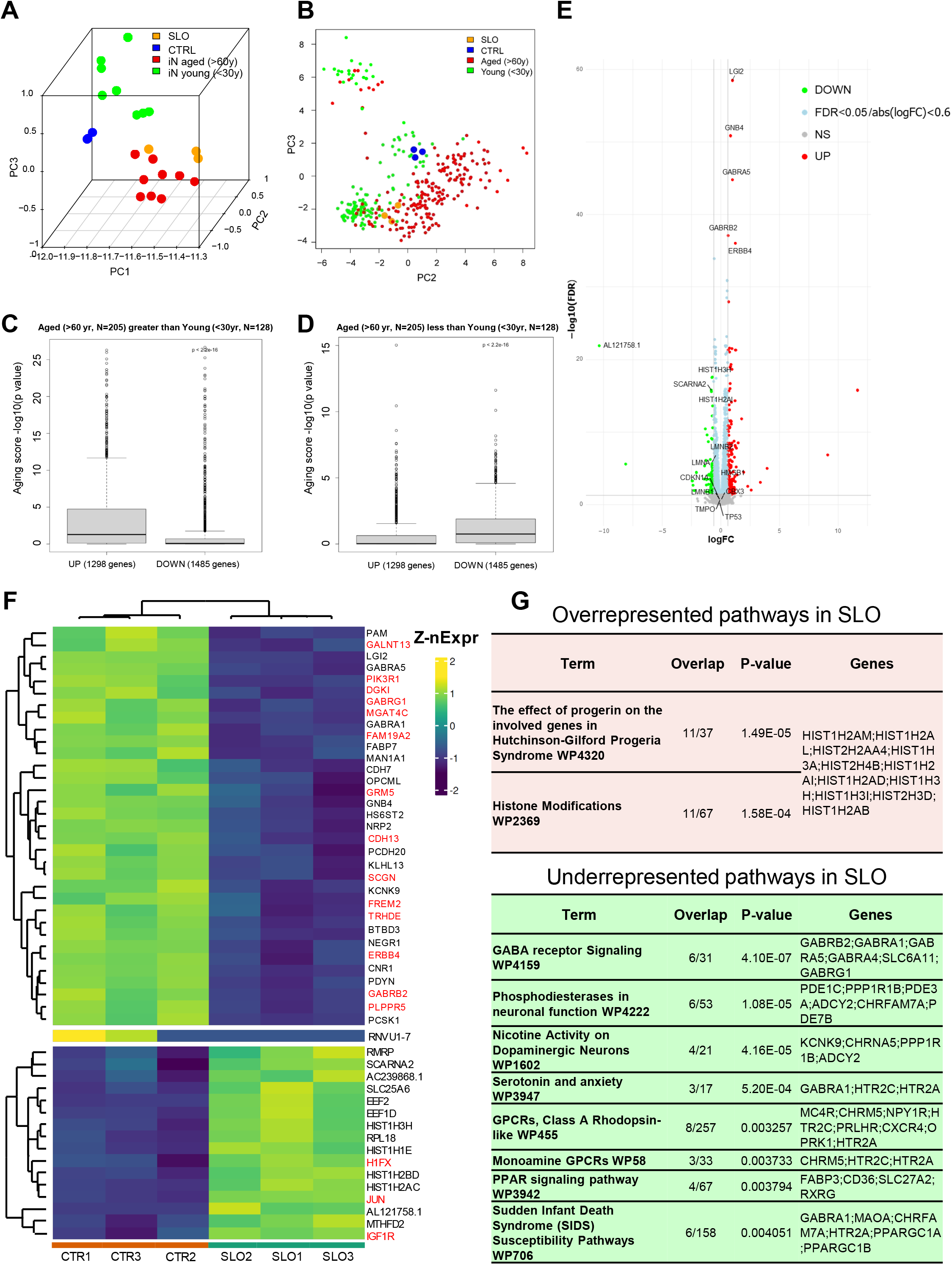
RNAseq analysis on SLO treated cortical neurons. PCA plot for SLO and CTRL samples as well as induced neurons (iNs) converted from both aged (9 samples) and young (8 samples) fibroblasts (A) and the aged and young PsychENCODE samples (the old group (>60 years, N=205) and the young group (<30 years, N=128)) (B). Boxplots for human aging scores association between SLO neurons and brain samples for up regulated genes (C) and down regulated genes (D) in the aged brains. Smear plot represents each gene with a dot, the gray dots (below cut off line) are genes with no change relative to the contrast direction, red and green dots denote up- and down-regulated expression, respectively, at an adjusted p-value (FDR) significance threshold of 0.05. The light blue dots are transcripts with FDR<0.05 but have log expression change of less than 0.6. The X-axis (log2 fold change) is the effect size, indicating how much expression has changed with SLO treatment (E). Heatmap clustering for 50 of the most differentially expressed genes with a p-value <0.05 and a log (2) fold-change greater or less than 2. The Z-score of given expression value is the number of standard-deviations away from the mean of all the expression values for that gene (F). All DEGs with a FDR <0.05 and 0.6≤logFC≤-0.6 are selected and tested for over- or under-representation of pathways in the gene list. Any significantly enriched WikiPathway pathways are ordered from most to least significant based on the p-value (G).

We then looked at the human aging scores (-log10(p-value) for the genes that are associated with aging, see RNAseq in methods) of cortical neurons treated with SLO. We found that the up-regulated genes in SLO-treated cells have higher human aging scores in the PsychENCODE aged group than the down-regulated genes (Figure 5C, t.test p<2.2e-16). Similarly, the down-regulated genes have significantly higher human aging scores in the PsychENCODE young group than the aged genes (Fig. 5D, t.test p<2.2e-16). These results suggest that our SLO-treated neurons have a similar gene expression dynamic to that in the aged human brain in PsychENCODE.

Comparison between SLO treated neurons and DMSO control neurons resulted in **271** differentially expressed genes (DEGs) (FDR < 0.05, 0.6≤logFC≤**-**0.6) with **190** genes down-regulated and **81** genes up-regulated upon SLO treatment (Figure 5E, Table S3). These DEGs are also present in the gene list that are significantly modulated by age in PsychENCODE (Table S4). In our SLO DEG list, GABA receptors are among the most down-regulated genes whereas histone variants are up-regulated (Figure 5F). Pathway analysis for DEGs in the SLO treated neurons revealed that neurotransmitter receptor signaling and GPCR signaling are down-regulated whereas pathways in the histone modification (especially histone variants) are up-regulated (Figure 5G).

Premature aging syndromes that are associated with mutations in LMNA or WRN genes resemble normal aging in terms of gene expression ^21, 22^. Over-expression of mutant Lamin A/C (Progerin) in normal neurons causes aging phenotypes ^7^. Interestingly, the SLO-treated neurons exhibited an upregulated pathway (WP4320) that shares 11 genes (30% of total genes in the pathway) involved in Hutchinson-Gilford Progeria Syndrome (Figure 5G). They included histone variants, several of which are involved in the histone modification pathway (WP2369). Other transcripts that are up regulated in SLO-treated cells included insulin receptor substrate 1 and 2 (IRS1 and IRS2), pro-apoptotic genes (FOXO3, BAD and BCL2L11), nutrients sensing transcripts (EIF4EBP1, TSC2, EEF2) and downstream kinase molecules (PIK3R2, ELK1, PTPRF, MAPK7, AKT1, MAP2K2, PLCG1, CRTC1 and JUN), and other transcripts (SHC2, RAB3A, DOCK3, RELA, NCK2, RACK1, SH2B1, LINGO1, STAT5B, EGR1, SQSTM1). Other down regulated transcripts in SLO treated cells included AMPA and NMDA receptors (GRIA1, GRIA2, GRIA3, GRIN2B), both trkB and trkC receptors (NTRK2, NTRK3) and their downstream calcium signaling molecules (NFATC4, CAMK2A, CAMK4), MAPK responsive transcripts (MAP2K1, KIDINS220, PRKAA2, PPP2CA) and other transcripts (GABRB3, MEF2C, SHC3, RASGRF1, PIK3R1, CDC42, CDH2, CNR1, SPP1, EIF4E, NSF, PTPN11, DLG1, APC). The transcriptome data suggest that the SLO-treated neurons resemble those from aged human cortex and premature aging samples.

### Induction of CS accelerates disease phenotype manifestation in ALS MNs

Neurodegenerative diseases such as amyotrophic lateral sclerosis (ALS) usually manifest symptoms after the 5th decade of life. We hypothesized that induction of CS in ALS iPSC-derived neurons would accelerate the presentation of disease phenotypes. We used the *TARDBP* mutant (298S) iPSC line generated from an ALS patient and its isogenic cell line (298G) produced by correcting the mutation using CRISPR for investigation of the ALS disease phenotype (Figure S4). Both mutant and corrected iPSCs were differentiated to spinal motor-neurons (MNs) using our previously published protocol ^3, 23^ (Figure 6A) and the MNs were treated with the SLO cocktail for 4 days. As expected, addition of the small molecules at day 25 and assayed at day 28 did not significantly alter the percentage of cells expressing cleaved caspase 3 (Figure 6B), indicating no obvious cytotoxicity for SLO and SSO treatments (control=26.9±1.3, SLO=25±2.4, SSO=24.7±4.9,) whereas we found significant number of caspase 3 positive cells in MG132 treated neurons (MG132=36.92±1.33).

**Figure 6.**
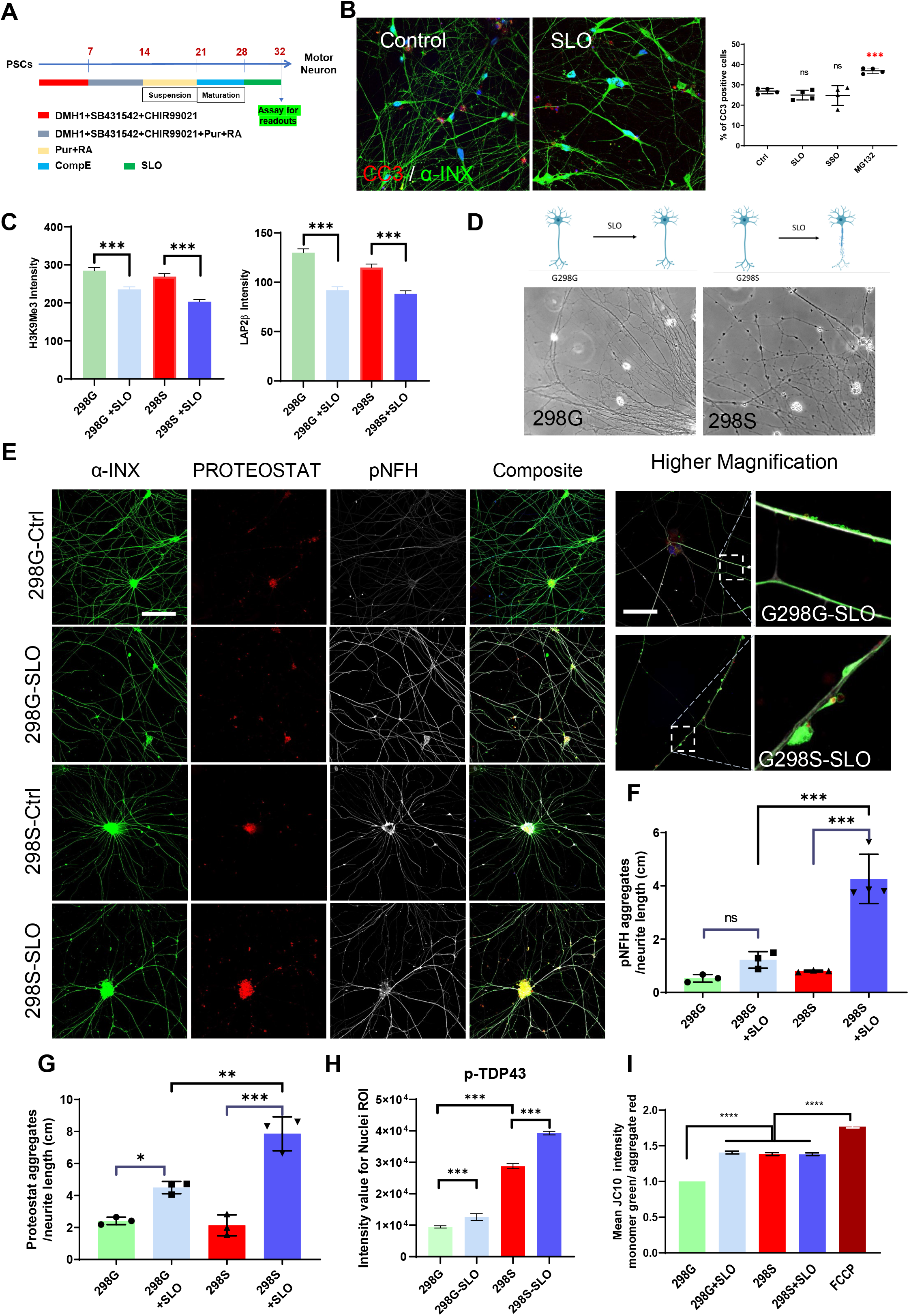
Phenotype presentation by SLO-treated motor-neurons derived from *TARDBP* mutant iPSCs. Differentiation protocol used for generating MNs from TDP-43 298S mutant and 298G isogenic iPSCs (A). Immunostaining for cleaved caspase 3 and alpha internexin proteins in cultured MNs treated with SLO, SSO and MG-132, at day 28 (B). High content imaging for H3K9M3 and Lap2β in both TDP43 298G isogenic control and 298S mutant following SLO treatment (Mean of SLO treatment compared to the control group with DMSO). Representative phase contrast image of MN cultures from both control and mutant ALS neurons treated with SLO (D). Immunostaining images for alpha-internexin, Proteostat, phosphorylated neurofilament heavy proteins in control and mutant MNs treated with SLO; right panel shows higher magnification images of control and mutant MNs treated with SLO (Scale bar=200 μm, for higher magnification images scale bar=50 μm) (E). Quantification result for phosphorylated neurofilament aggregates (F) and Proteostat positive protein aggregations (G) and phosphorylated TDP43 protein (H) across all groups. Mitochondrial membrane potential (JC10 assay) evaluation of ALS-iPSC derived MNs treated with SLO compared to the healthy isogenic control cells and isogenic cells treated with FCCP (I). (ns: not significant, *: p<0.05, **: p<0.01, ***: p<0.001 one-way ANOVA with Dunnett’s multiple comparison test, for JC10 assay data was quantified using 15,000 cells per group from two independent experiments. Statistical analysis was performed using One-way ANOVA, Tukey post-hoc test (****-p<0.001)).

We then examined if the MNs treated with SLO display CS-like phenotypes as we observed in fibroblasts and cortical neurons. Both 298G and 298S MNs showed a reduction in the expression of H3K9Me3 and Lap2β following SLO treatment (Figure 6C), indicating that SLO treatment induces CS. Both cell lines responded at the same level to the SLO chemicals and difference in H3K9Me3 and lap2β signal intensity was not significant (Figure 6C). We then asked if induction of CS accelerates neuronal degeneration. By day 32, the 298S MNs treated with SLO showed axonal swellings, a sign of axonal degeneration whereas 298G MNs showed relatively intact neurites (Figure 6D). Proteostat staining was increased in SLO-treated cells, especially in the 298S MNs. ALS MNs, when treated with SLO, were positive for phosphor-TDP43 and Proteostat (Figure 6E, S5A). Immunofluorescence for phosphorylated neurofilament, a marker for axonal degeneration and turnover, was also significantly increased in the SLO-treated 298S than in non-SLO-treated 298S and SLO-treated 298G MNs. Under higher magnification Proteostat positive aggregates were observed along the axons and were positive for both α-internexin and phosphorylated neurofilament (Figure 6E, S5A). Quantification of the aggregates showed a significant increase in p-NFH aggregates (1.22 ±0.31 in 298G compare to 4.26±0.92 in 298S) and Proteostat-positive aggregates (4.49±0.38 in G298G compare to 7.86±1.06 in G298S) in 298S ALS mutant MNs than the 298G isogenic control MNs that were treated with SLO (Figure 6F, G). Following SLO treatment we detected TDP43 signal permeation from nucleus to the cytoplasm area (Figure S5A), and ALS MNs treated with SLO had significantly more p-TDP43 signal compared to the isogenic control and cells that are not treated with SLO (Figure 6H, S5A). Interestingly, neurite swelling contained p-TDP43 proteins that are co-labeled with Proteostat and other neurofilaments (Figure S5A and 6E).

One of the hallmarks of the neurodegeneration is mitochondrial deficit. Assay with JC-10 staining showed that ALS-MNs had a lower mitochondrial membrane potential (MMP) than the isogenic control MNs. Treatment with SLO lowered the membrane potential for control neurons but not further for the ALS-MN (Fig 6I). Morphological analysis with Mitotracker staining showed that isogenic control MNs treated with SLO had shorter and smaller mitochondria than those without SLO treatment, but SLO treatment had no further effect on ALS MNs (Figure S5B,C). Giving that mitochondrial DNA is generally not reprogrammed during iPSC generation, these results indicate that the mitochondrial phenotypes are present in ALS-iPSC-derived MNs and SLO treatment does not change the mitochondrial phenotypes beyond what it was in ALS-MNs, suggesting that senescence- or disease-related mitochondrial phenotypes are retained during reprogramming.

### Autophagy Induction clears up protein aggregation and improves neurite health

The fast and consistent presentation of disease relevant phenotypes in SLO-induced cultures makes them amenable for testing therapeutic agents. We examined the effects of molecules on protein aggregation and neurite swelling in the SLO-treated ALS motor neuron cultures, including the current ALS medication Edaravone, autophagy activators STF-62247, SMER28, Flubendazole, and the peptide Tat-Beclin, and KU-60019, a molecule identified from our initial cell toxicity screening as neuroprotective. In addition, Amiodarone was used as an unrelated hit. MNs from both ALS (298S) and isogenic (298G) iPSCs at day 28 post differentiation were treated with SLO and then the compounds were added separately 24 hours later, and cells were analyzed at day 32 (Figure 7A). SMER28 and Tat-Beclin decreased proteostat positive aggregates in both ALS (37%±14 and 62.5%±5.5) and isogenic MNs (38%±11.7, 66.9%±5 for 298G) as compared to SLO treated control. Edaravone and KU-60019 reduced the level of Proteostat, more so on ALS cells (to 26%±6 and 17%±0.7) than the isogenic cells (64%±10.4, 30%±13.6 for 298G). MNs treated with STF-62247 and Flubendazole showed more aggregates in 298G cells compared to the SLO treated cells (179%±29, 143±26.8) and no improvement in 298S cells. Amiodarone did not improve protein aggregation (Figure 7B).

**Figure 7.**
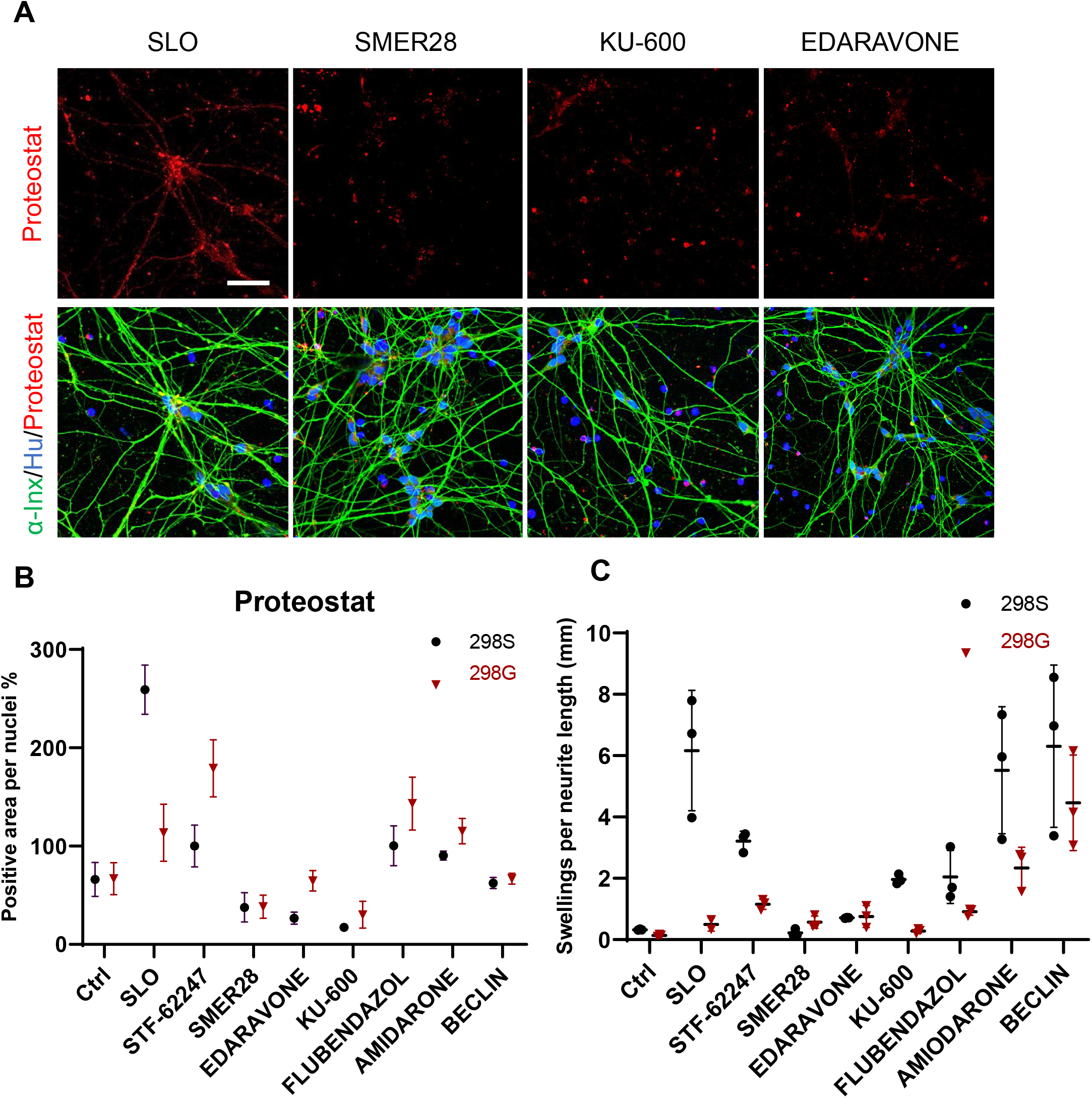
Testing molecules that rescue the disease phenotype in ALS MNs. MNs from TDP-43 G298S mutant and G298G isogenic iPSCs treated with SLO to induce CS phenotypes and 24hr later candidate molecules SMER28, EDARAVONE and KU-60019 were added to the culture and cells stained with Proteostat dye for protein aggregation and alpha-Internexin for visualizing neurites (A). Quantification of immunostaining images positive for Proteostat (B) and neurite swellings (C). (Scale bar=100 μm)

Neurite swelling is one of the obvious morphological changes in ALS MNs following SLO treatment. SMER28 and EDARAVONE significantly reduced the number of swellings to the level of isogenic control cells (Figure 7C). However, KU-60019 treatment did not improved the swelling phenotype to the normal level (Figure 7C) despite significant reduction in protein aggregation (Figure 7B). Other molecules did not show significant improvement in 298S MNs or even induced more swellings in the 298G control cells (Figure 7C). Thus, EDARAVONE and SMER28 can reduce both protein aggregation and neurite swellings in 298S TDP43 mutant cells and were beneficial for MN health in our CS culture system.

## Discussion

Most neurodegenerative diseases are concurrent with aging ^24, 25, 26, 27^. Hence, recapitulating CS in stem cell derived neurons could expand the capacity of the iPSC model to study disease mechanisms ^28, 29^. By using H3K9Me3, HP1γ and Lap2β as readouts and screening for chemicals/pathways that induce CS in neonatal fibroblasts and iPSC derived cortical neurons, we developed cocktails of small molecules that induce CS in the cortical neurons. This chemically induced senescence (CIS) approach was validated in motor neurons derived from ALS patient iPSCs. Importantly, CIS enhanced the presentation of disease related phenotypes. This CIS strategy will likely enable more effective iPSC-based modeling of age-related degenerative diseases and enable better therapeutic target design.

Cellular senescence across different cell types shares different features including mitochondrial dysfunction, DNA damage, P16 expression changes and epigenetic marks for gene silencing ^19, 30, 31, 32^. These alterations ultimately result in age-related changes at the cellular level, including changes in cell size, shape and metabolism, proliferation arrest, and telomere erosion ^15, 33, 34^. In mitotic cells like fibroblasts, expression of P16 accompanies proliferation arrest and induces senescence ^35, 36^. P16 activation by Palbociclib in our study is one of the most efficient pathways in CS by blocking CDK4/6 and proliferation of fibroblasts, causing senescence. Other pathways in our study with fibroblasts are related to the DNA repair, DNA synthesis and DNA alkylation pathways; all related to cell division and telomere attrition. Surprisingly none of the sirtuin inhibitors induced senescence in fibroblasts or neurons despite their effects on aging ^32, 37^. This may reflect differences between cell types or insufficient treatment with inhibitors.

In post-mitotic cells like neurons, protein quality control, including proteasome and autophagy processing, is more important in CS progression ^38, 39, 40^. This is reflected in our study showing the powerful CS-inducing effect of autophagy inhibitors (SBI-0206965). Faulty autophagosomes could not clear impaired mitochondria and unfolded protein debris, leading to lack of mitochondrial turnover and producing more oxidative stress ^41, 42^. Oxidative stress generates ROS and accounts for higher DNA mutations, which is ultimately related to CS ^16, 43^. Similarly, we found that inhibition of DNA glycosylase (*OGG1*), important in detecting and removing oxidized nucleotides in genomic DNA, exacerbates CS phenotype in neurons but not in fibroblasts. Two of three small molecules in SLO, DNA glycosylase inhibitor (O151) and HIV protease inhibitor (Lopinavir), modulate CS phenotypes in neurons, indicating that base excision repair (BER) pathway is critical for neuronal health and is linked to neurodegenerative diseases ^44, 45^. Lopinavir also inhibits ZMPSTE24 ^46, 47^, thereby blocking lamin A biogenesis and leading to an accumulation of prelamin A. ZMPSTE24 deficiency in humans causes an accumulation of prelamin A and leads to lipodystrophy and premature aging ^48, 49, 50^, which perhaps causes senescence phenotype in our cultured neurons. We used three different endopeptidase inhibitors Phosphoramidone (a general metalloendopeptidase), Lopinavir (zinc metalloprotease inhibitor) and GGTI-298 (a geranylgeranyltransferase I (GGTase I) inhibitor), and only Lopinavir induced senescence in cortical neurons in all three markers. Interestingly, none of the endopeptidase inhibitors induced senescence phenotype in fibroblasts, indicating that neurons are more sensitive to the activity of endopeptidase, perhaps for the processing of other lamin proteins rather than just for Lamin A ^51, 52^.

Information on CS derives largely from studies on mitotic cells. Transcriptomic analysis revealed that the gene expression pattern of our SLO-treated neurons resembles that of iNs and aged brain. In particular, our in-vitro neuronal senescence system, despite the lack of many other cell types that are normally present in the human brain, resembles the aging cortex samples as indicated by the substantial overlap of age-related transcripts between our CIS neurons and aged human brain tissues ^20^. These transcription profiles may be more specific to CS in neurons. For example, transcripts that are involved in neurexin/neuroligin complexes at synaptic membrane assembly and neurotransmitter release from GABA, glutamate and cholinergic systems are common between aged brains and SLO induced senescence. Neurexin expression declines with age and causes decrease in synaptic density and cognitive decline ^53, 54^. Other transcripts like CREB Regulated Transcription Coactivator 1 (CRTC1) transcription coactivator of CREB1 ^55^, which show significant change in our SLO-treated cortical neurons, also contribute to brain aging and neuronal senescence. Some of other molecules such as p62 (SQSTM1) have multiple function and contribute to neurodegeneration by binding to the ubiquitin molecules that are marked for degradation and by binding to autophagy molecule LC3-II ^56^. In addition, our CIS neurons share several histone variants with the progerin effect in the progeria syndrome. Histone variants are one of the most affected transcripts during CIS in the cortical neurons. Histone variant exchange, by regulating expression of age related genes ^57^ and/or chromatin organization ^58^, is one of the mechanisms behind CS and aging. Thus, our CIS recapitulates aspects of premature aging effects primarily at the epigenetic level.

A major driving force behind the development of CIS is to enable effective and reliable modeling of age-related diseases using human stem cells. We and others have shown that some aspects of neurodegenerative changes such as ALS may be recapitulated by strictly controlling the neuronal differentiation process, prolonged maturation, and undergoing stress (including culturing under a basal condition without trophic factors and medium changes) ^3, 4, 59^. Such manipulations over a long term adds variables to the system, making stem cell-based disease modeling more difficult. MNs from patients with TARDBP mutations have increased levels of soluble and detergent-resistant TDP-43 and show decreased cell survival, suggesting that this model is representative for ALS pathology ^60, 61^. However, neither increase in insoluble TDP43 nor its mis-localization phenotype is repeated in a recent study 62. Similarly, dopamine neurons from Parkinson’s disease (PD) iPSCs exhibited mitochondrial dysfunction and oxidative stress, changes in neurite growth and morphology, synaptic connectivity and lysosomal dysfunction ^63, 64, 65, 66^, but hallmark pathology like protein aggregation and Lewy body formation are rarely observed ^64, 65, 66, 67^. These inconsistencies may be due to the different protocols used and the long-term cultures that are necessary to mature the stem cell derived neurons. The current CIS approach enables an early and consistent presentation of disease relevant phenotypes, including protein aggregation and axonal degeneration in TDP43 mutant MNs. Since the cocktails induce CS in different neuronal types, it is likely that the CIS approach may promote phenotypic presentation in other age-related diseases using iPSCs.

Our CIS method induces CS in a short period (after 2-4 days of treatment) without a need of genetic manipulation. It promotes reliable and consistent presentation of disease relevant phenotypes and it is not specific to any particular disease. The cocktails were developed by screening a relatively small pool of molecules, suggesting that other molecules, especially those affecting similar pathways, may also induce CS. Since our CIS method enables faster and consistent presentation of disease phenotypes from iPSC-derived neurons, it is also useful for establishment of drug testing platforms. As a proof of principle test, we found that the current ALS medication Edaravone and one of the many autophagy activators SMER28 but not others mitigate protein aggregation and neurite swelling in ALS iPSC-derived motor neurons, highlighting the utility of the system.

## Supporting information

Table S2- Small molecules response

Table S3- DEG

Table S4 SLO vs Brain Samples

## Acknowledgements

We thank D. Moore lab for providing EtO and XTPNgn2: 2A:Ascl1 (N2A) vectors. We are grateful for technical support provided by K.M. Knobel and CMN Core at Waisman center. We would also like to thank R. Bradly, J.R. Jones, Y. Tao, M. Ayala for their comments and technical support. This study was supported in part by the ALS Association (20-IIP-556), the NIH-NIMH (MH100031), NIH-NINDS (NS086604, NS096282), NIH-NICHD (HD076892) and a core grant to the Waisman Center from the NIH-NICHD (U54 HD090256). S-C.Z. acknowledges the Steenbock professorship in Behavioral and Neural Sciences.

## Author Contributions

A.F.: Design and conception of the study, writing of manuscript, maintenance, directed differentiation, direct conversion of fibroblasts, establishing of high content imaging assays, small molecule screen. S.M: conduct the SLO response in cortical neurons, western blotting and JC10 MMP assay, writing of manuscript. L.K: Mitotracker assay and MMP assay. A.J.P.: gene targeting of TARDBP in human PS cells and characterization of iPSCs. C.R.K.H.: Immunostaining, high content imaging, data interpretation, editing the manuscript. J.B and J.M.M.: RNA sample preparations, immunostaining and cell toxicity assays. A.B.: design and interpretation of small molecule screen and follow-up experiment, writing of manuscript. D.W: RNAseq data analysis, comparing RNAseq data to iNs and brain data, writing of manuscript. S-C.Z.: design and conception of the study, data interpretation, writing of manuscript.

## Resource Availability

### Lead Contact

Further information and requests for sources should be directed to, and will be fulfilled by, the Lead Contact, Dr. Su-Chun Zhang.

### Materials Availability

Plasmids and hESC/hiPSC lines generated in this study are available from the Lead Contact with a completed Materials Transfer Agreement.

### Data and Code Availability

The published article includes all datasets generated or analyzed during this study. The datasets supporting this study are available from the lead contact, Dr. Su-Chun Zhang upon request.

## Experimental Model and Subject Details

### Neuronal differentiation from hPSCs

Human embryonic stem cells (H9 or WA09, WiCell), H9-GFP (AAVS1-CAG-eGFP) cells and TARDBP mutant (G298S) and isogenic (control) induced pluripotent stem cells (iPSCs) were grown on Matrigel with Essential-8 medium (Stemcell Technologies) to 25% confluency. For cortical neuron differentiation, the fifth day cultures of hPSCs were treated with Accutase and the dissociated single cells were cultured in the neural differentiation medium (NDM) (DMEM/F12:Neurobasal 1:1 + 1X N2 Supplement + 1mM L-Glutamax) with the SMAD inhibitors SB431542 (Stemgent), DMH-1 (Tocris) (both at 2μM) and Rho kinase inhibitor (Tocris) (overnight) as spheres (embryoid bodies) for seven days. On day 8, neural spheres were patterned to dorsal forebrain (cerebral cortical) progenitors with the smoothened antagonist cyclopamine (Stemgent, 2μM) and FGF2 (R&D, 10 ng/ml) for seven days. On day 14, neural progenitors were dissociated with Accutase to single cells and plated on Laminin coated plates in the maturation media (DMEM/F12/Neurobasal 50%/50%, 1x B27 Supplement, 1x Non-essential amino acids, 1x Glutamax) supplemented with Compound E (0.1 μM, TOCRIS) for final maturation until assay time. For motor neuron differentiation, we used our previous published protocol with no further modification 23. For SLO experiments and autophagy activation MNs treated four days after maturation (Day 25) with SLO molecules and autophagy activators and other molecules added 24hr later and neurons cultured for another 3 days and analyzed at day 29.

### Direct Conversion of Human Fibroblasts into iNs

Primary human dermal fibroblasts (WC-04-05-CO-DG, 72 year-old male, WC-60-07-CO-CMN neonatal male, WC-03-06-CO-DG, 62 year-old female, and WC-59-07-CO-CMN, neonatal female, neonatal fibroblasts from WiCell and aged fibroblasts from David Gamm’s lab), were cultured in DMEM containing 15% tetracycline-free fetal bovine serum and 0.1% NEAA (Life Technologies), transduced with lentiviral particles for EtO and XTPNgn2: 2A:Ascl1 (N2A), and expanded in the presence of G418 (200 mg/ml; Life Technologies) and puromycin (1 mg/ml; Sigma Aldrich). For iN conversion, we followed the previously published protocol 13. We used neuron conversion (NC) medium based on DMEM:F12/Neurobasal (1:1) for 3 weeks. NC contained the following supplements: N2 supplement, B27 supplement (both 1x; GIBCO), doxycycline (2 μg/ml; Sigma Aldrich), Laminin 1 μg/ml; (Life Technologies), dibutyryl cAMP (400 μg/ml; Sigma Aldrich), human recombinant Noggin (150 ng/ml; R&D), LDN-193189 (0.5 μM; Cayman Chemicals) and A83-1 (0.5 μM; Stemgent), CHIR99021 (3 μM; LC Laboratories) and SB-431542 (10 μM; Cayman Chemicals). Medium was changed every third day. For further maturation, iNs were cultured in DMEM:F12/Neurobasal-based neural maturation media (NM) containing N2, B27, GDNF, BDNF (both 20 ng/ml; R&D), dibutyryl cAMP (400 μg/ml; Sigma Aldrich), doxycycline (2 μg/ml; Sigma-Aldrich), and laminin (1 μg/ml; Life Technologies). Converted neurons in 96 well plates were used for immunostaining without further purification.

### Immunofluorescent staining and microscopy

Cells were fixed for 20 minutes with 4% paraformaldehyde in PBS at a room temperature. Samples were blocked with 4% donkey serum and 0.2% Tween20 for one hour. Primary antibodies were diluted in 4% donkey serum and 0.1% Tween20 and applied to samples overnight at 4°C. Samples were washed with PBS, incubated with fluorescein-conjugated secondary antibodies for one hour at room temperature, and counterstained with Hoechst for 20 minutes. Samples were imaged on a Nikon A1s confocal microscope (Nikon). For measuring neurite length and swellings, images were processed with Fiji software. First, a threshold was set for images to select all cell processes, then neurites were skeletonized. For analyzing the skeletonized neurites prune cycle method were used and parameters set to the shortest branch and end points eliminated to prune ends. Then total brunch length was calculated for labeled skeletons (total branch length in pixel/10,000=branch length in cm) and for aggregates per neurites length total number of aggregates were divided by the branch length.

The following primary antibodies were used:

| Antibody | Species | Catalog No. | Company | Dilution |
| --- | --- | --- | --- | --- |
| TDP43 | Rabbit | 10782-2-AP | Proteintech Group, Inc. | 1000 |
| phospho-TDP43 | Rat | MABN14 | MilliporeSigma | 500 |
| P-SCG10 | Rabbit | STJ91270 | St. John's Lab | 300 |
| H3K9Me3 antibody | Rabbit | ab176916 | abcam | 5000 |
| CHAT antibody | Goat | AB144P | Chemicon | 1000 |
| HB9 | Mouse | 81.5C10 | DSHB | 50 |
| TUBB3 | Rabbit | PRB-435P | Covance | 10,000 |
| MAP2 | Mouse | M1406 | Sigma | 1000 |
| Lamp-2 | Mouse | NBP2-22217 | Novus Biologicals | 250 |
| Lamin B2 antibody | Mouse | ab8983 | abcam | 500 |
| HP1γ antibody | Rabbit | 2619 | Cell Signaling | 1000 |
| HP1γ antibody | Mouse | MABE656 | Millipore | 500 |
| LAP2β | Mouse | 611000 | BD Biosciences | 50 |
| Lamin A + C antibody | Rabbit | ab133256 | abcam | 500 |
| H2A.X | Mouse | 05-636 | Upstate (EMD Millipore) | 500 |
| Alpha-Internexin | Rabbit | AB40758 | abcam | 500 |
| Cleaved Caspase-3 | Rabbit | 9661S | Cell signaling | 300 |
| PhosphoDetect™<br>Anti- Neurofilament H | Mouse | NE1022 | Millipore-Sigma | 2000 |
| Proteostat | NA | ENZ-51023 | Enzo | 1000 |
| SOX1 | Goat | AF3369 | R&D systems | 2000 |
| OTX2 | Goat | AF1979 | R&D systems | 2000 |

### High-content imaging

For measuring cell population, fluorescence intensity, apoptosis, and intensity of H3K9Me3, Lamin B2, Lap2β, and HP1γ, cells were plated in 96-well imaging plates (18000 cells per well, CELL CARRIER) and treated with different molecules (supplementary table S1). After staining, images were analyzed using the high-content cellular analysis system Operetta (Perkin Elmer). A set of 60 fields was collected from each well (total of three wells per treatment) using the 40× objective, resulting in over 10,000 cells being scored per well. In our analysis workflow, we first identified the nuclei based on default protocol B and calculated the intensity and morphology properties for each nucleus by gating out nuclei with a roundness of below 0.75 and intensities above 1500 for removing extremely bright nuclei (dead cells). We then calculated the signal intensity for each protein in different channels separately. For quantification of H3K9Me3, Lamin B2, Lap2β intensity in directly reprogrammed neurons, we identified the cytoplasm based on the βIII-tubulin staining surrounding each selected nucleus and quantified the expression of markers in βIII-tubulin positive population. All raw data were exported and analyzed with GraphPad Prism (GraphPad Software).

### RNA-seq procedure

Cortical neurons differentiated for 7 days and then treated with SLO for 5 days were collected for RNAseq analysis. All experiments were run three times and RNA was extracted from all samples (3 biological replicates and 3 technical replicates) using the RNeasy Plus Mini kit (Qiagen) following manufacturer’s instructions. RNA quality was assessed using an Agilent RNA PicoChip with all samples passing QC. Sample libraries were prepared using poly-A selection using an Illumina TruSeq RNAv2 kit following manufacturer’s instructions. Prepared libraries were sequenced for 101-bp single-read and performed on an Illumina HiSeq to a read depth of >25 million reads per sample by the DNA Sequencing Facility in the University of Wisconsin-Madison Biotechnology Center. FastQC was performed on all samples with every sample passing all quality control measurements.

### RNA-seq analysis

Differentially expressed genes were identified with a glm function using the edgeR package. A subset of up to 50 of the most differentially expressed genes with a p-value <0.05 and a log fold-change greater or less than +/− 2 were selected according to their FDR rank in the list of DEGs23. Next, both samples and genes were clustered using Euclidean distances. For genes, an additional elbow function was applied to estimate the number of gene clusters present. Calculated relationships are depicted by dendrograms drawn at the top (samples) and to the left (genes) of the heatmap. The gradation of color is determined by a Z-score that is computed and scaled across rows of genes normalized by TMM. The Z-score of a given expression value is the number of standard-deviations away from the mean of all the expression values for that gene.

The empirical Bayes hierarchical modeling approach EBSeq was used to identify differentially expressed genes across 2 or more conditions. Median normalization technique of DESeq was used to account for differences in sequencing depth. EBSeq calculates the posterior probability (PP) of a gene being in each expression pattern. Genes were declared differentially expressed at a false discovery rate controlled at 100*(1-α) % if the posterior probability of P1 (EE) is less than 1-α. Given this list of DE genes, the genes are further classified into each pattern and sorted by PP.

For quantifying the degrees to which genes are associated with aging in the human brain, we performed one-side t-tests for each gene to determine if it is significantly positively expressed (i.e., up-regulation) in the aged group (>60 years, N=205) and the young group (<30 years, N=128) of the healthy human brain tissue samples (DLPFC) in the PsychENCODE project ^20^. Then, we used the value of -log10 (t-test p value) of the gene as “human aging score” to quantify its degree of association with the corresponding group in the human brain. Finally, each gene has two human aging scores to quantify its association with (1) up-regulation in the aged group; (2) up-regulation in the young group.

### Principal component analysis

For principal component analysis (PCA), all the data including cortical neurons RNAseq data, iN data and PsychENCODE data were first transformed by log10 (x+1). All samples including our SLO and CTRL samples, the aged and young groups in PsychENCODE (as described above), and iNs. combined as a single data matrix (samples by genes) for PCA.

### Pathway Analysis

DEGs from each group were analyzed for differentially regulated pathways using ENRICHR (www.enrichr.org) which utilizes several pathway databases for general pathway analysis. For our analysis, the KEGG and Wikipathway databases were utilized. DEGs were defined as >100 TPM and >2-fold change over each of the other groups. Pathways that were statistically significant were highlighted as potential differentially regulated. Only pathways that were found significant in more than one of the three analyses were considered for further evaluation.

### qRT-PCR

RNA samples were obtained using the RNeasy Plus Mini kit (Qiagen) following manufacturer’s instructions. cDNA libraries were constructed using iScript cDNA Synthesis kit (Bio-Rad) using 500ng of purified RNA from each sample as input following manufacturer’s instructions. qRT-PCR was performed on a CFX Connect qPCR machine (Bio-Rad) using iTaq SYBR green supermix (Bio-Rad) and equal amounts of cDNA samples. Results were normalized to GAPDH or 18s rRNA levels using the ΔΔCt method.

### SA-β Galactosidase assay

Fibroblasts were fixed using 1X fixation buffer provided in reagents and procedure were performed following manufacturer’s instructions for Cellular Senescence Assay Kit (Sigma, KAA002). Bright-field mages were acquired using a Nikon microscope and positive cell numbers calculated using the Fiji software. Positive cells were grouped based on their appearance after β-Gal staining using histogram function (quantity of staining) to the high and moderate.

### Live and Dead cell staining

For the cell toxicity assay, cells were plated in 96 well optical plates at a density of 30,000 cells per well and each 3 well (experimental replicates) treated with different small molecules for 24 hr. Then cells were washed with PBS and incubate with 1 μM EthD-1 and 1 μM calcein AM in the LIVE/DEAD™ Viability/Cytotoxicity Kit (Thermo Fisher, L3224) for 30 min at RT and imaged using Operetta (Perkin Elmer) and analyzed with Harmony software.

### Single nucleotide polymorphism (SNP) modification in TARDBP locus

To perform single nucleotide polymorphism (SNP) modification, we utilized the single-strand oligonucleotide (ssODN) method. Following sgRNA identification for the site of interest using the crispr.mit.edu design tool, we cloned the sgRNA sequences into the pLentiCRISPR-V1 plasmid from the laboratory of Feng Zhang (not available through Addgene anymore, but V2 version is plasmid #52961) following the protocol provided with the plasmid (Sanjana NE et al., 2014). Cells were cultured and electroporated as described in Chen Y et al., 2015. Single hESCs (1×10^7^ cells) were electroporated with appropriate combination of plasmids in 500 microliters of Electroporation Buffer (KCl 5mM, MgCl2 5mM, HEPES 15mM, Na2HPO4 102.94mM, NaH2PO4 47.06mM, PH=7.2) using the Gene Pulser Xcell System (Bio-Rad) at 250 V, 500μF in a 0.4 cm cuvettes (Phenix Research Products). Cells were electroporated in a cocktail of 15 micrograms of the pLentiCRISPRV1-TDP43 sg14 plasmid and 100 microliters of a 10 micromolar ssODN targeting the TDP43 G298S mutant genetic site. This ssODN was non-complementary to the sgRNA sequence and consisted of 141 nucleotides – 70 nucleotides upstream and 70 nucleotides downstream of the targeted indel generation site (Yang et al. 2013). Following electroporation, cells were plated on MEF feeders in 1.0 μM ROCK inhibitor. At 24- and 72-hours post-electroporation, cells were treated with puromycin (0.5 μg/ml, Invitrogen, ant-pr-1) to select for cells containing the pLentiCRISPRV1-TDP43 sg14 plasmid. After removal of the puromycin at 96 hours, cells were cultured in MEF-conditioned hPSC media until colonies were visible.

For genotyping single-cell generated colonies were manually selected and mechanically disaggregated. Genomic DNA was amplified using Q5 polymerase-based PCR (NEB) and proper clones determined using sanger sequencing. To identify non-specific genome editing, we analyzed suspected off-target sites for genome modification, using the 5 highest-likelihood off target sites predicted by the crispr.mit.edu algorithms.

### Mitochondrial morphology (Mitotracker) and membrane potential assay (JC-10 assay)

Neuronal progenitors were plated on Cellvis 35mm glass bottom dishes at 30,000 cells per dish and matured for 7 days. Mitotracker red (M7512, Invitrogen) were added directly to the culture media at final concentrations of 50nM and incubated for 15min in the incubator. Cells were then washed three times with phenol free neurobasal media (pre-warmed to 37°C) and switched to 2mL of pre-warmed neuronal media. Imaging was performed on a Nikon A1s confocal microscope with live cell chamber incubation. Nikon Elements software were used to acquire images under resonant scanning mode.

Mitochondrial membrane potential assay was performed using JC-10 mitochondrial membrane potential assay kit (ab112134, Abcam). H9 derived Cortical neuron progenitors and 298G and 298S hiPSC derived motor neuron progenitors were plated at 18000 cells per well in a 96 well imaging plate (Cell carrier). Neurons were treated with SLO (1:1000) at day 7 for cortical neurons and at day 4 for motor neurons. JC-10 assay was performed at Day 11 for cortical neurons and Day 9 for motor neurons. FCCP (mitochondrial uncoupler) at 2μM was used as a positive control. Neurons for positive control were treated with FCCP for 30 mins at 37°C followed by a wash with the complete neuronal medium. First, JC10 buffer A was added to the neurons and incubated for 30 minutes at 37°C. Then both JC10 buffer B and Nucblue live ready reagent (Thermo fisher) were added to the neurons and imaged immediately. Live cell imaging was acquired using High content microscopy (Operetta- Perkin Elmer). Image analysis was performed using Columbus software. Statistical analysis was made using GraphPad Prism 9.0.

### MitoSoX staining

Mitosox imaging assay was performed using MitoSOXTM Red Mitochondrial Superoxide Indicator purchased from Thermo Fisher (M36008). MitoSOX red (5μM) was added to the neurons and incubated for 30 minutes at 37°C. MitoSOX was removed after 30 minutes and Nucblue live ready reagent was added to the neurons and imaged immediately.

### Immuno blotting

H9 derived cortical neurons were gently scraped off the wells, washed with PBS and centrifuged at 1600 rpm for 2 mins. Pellets were lysed on ice using RIPA lysis buffer (Cell Signaling) supplemented with Halt Protease and Phosphatase inhibitor cocktail (Thermo Fisher Scientific) and 4-(2-Aminoethyl)benzenesulfonyl fluoride hydrochloride (AEBSF, Sigma). Samples were centrifuged at 18000rpm for 20 min at 4oC. Total protein concentrations were measured using Pierce BCA protein assay (Thermo Fisher Scientific). 2x Laemmli sample buffer (Bio-rad) was added to the protein sample and boiled at 95°C for 5 minutes. Protein samples (10μg/group) were run on 4-20% Mini-Protean TGX precast gel (Bio-rad), transferred to polyvinylidene difluoride (PVDF) membranes, blocked with 5% non-fat dry milk and then incubated with primary antibodies overnight at 40C. Signals were visualized using horseradish peroxidase conjugated secondary antibodies and captured with ChemiDoc system. The following primary antibodies were used: LAP2 (1:5000, BD Biosciences), H3K9Me3 (1:5000, Abcam), HP1ϒ (1:1000, Cell signaling).

