## Supplementary information for "Chemically Induced Senescence in Human Stem Cell-Derived Neurons Promotes Phenotypic Presentation of Neurodegeneration"

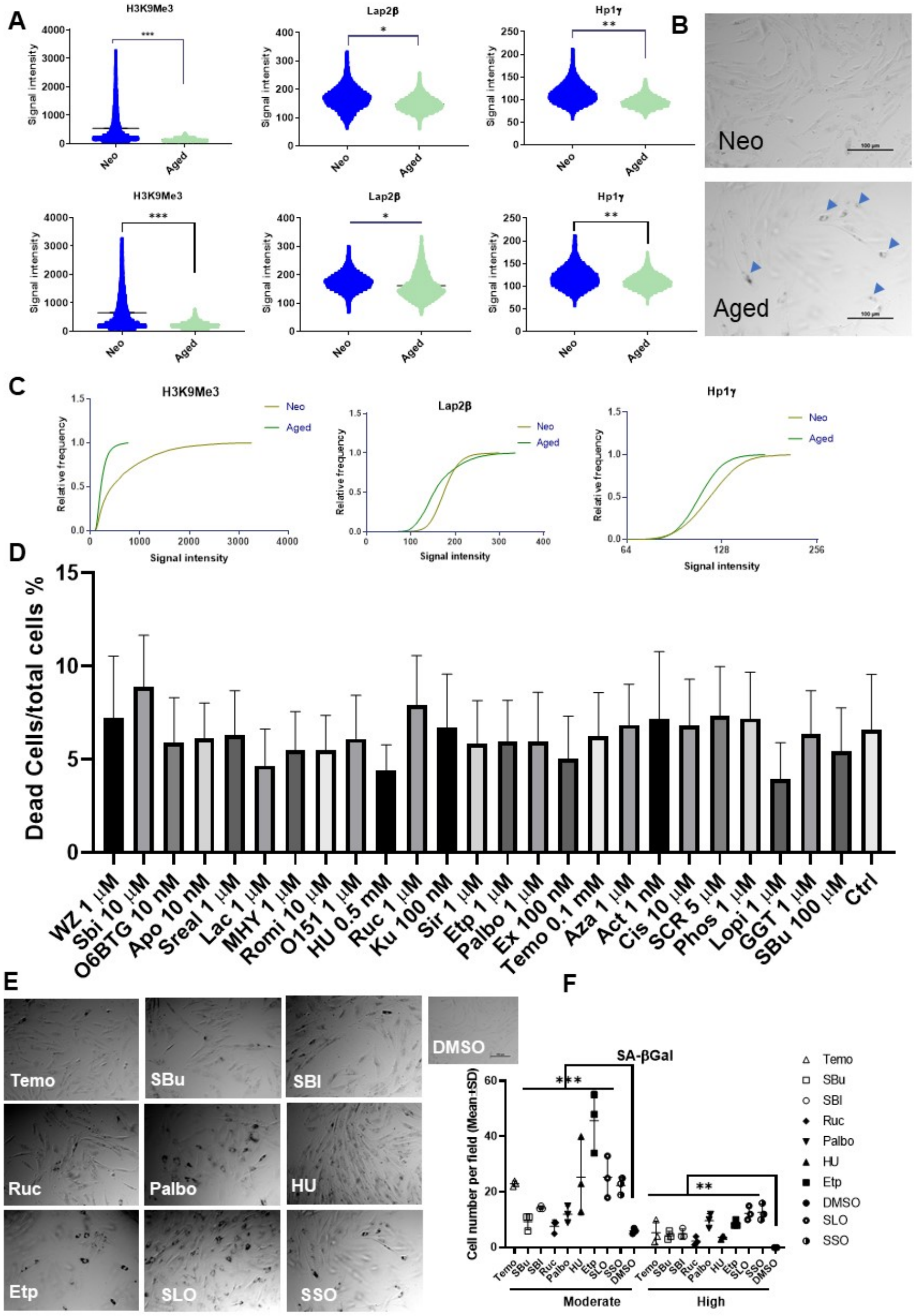


**Figure S1. Related to figure 1**, Individual values for H3K9Me3, Lap2β and HP1γ expression in both male (upper panel) and female (lower panel) fibroblast cells (A) and phase contrast images of senescence associated β-Galactosidase staining (arrowheads) for both neonatal and aged (female 62 years old) fibroblasts (B). Frequency distribution analysis of results from high content imaging for H3K9Me3, Lap2β and HP1γ proteins in female neonatal and aged (62 years old) fibroblasts (C). Cell toxicity assay for small molecules in the used concentration in actual experiment compared to the DMSO control in neonatal fibroblasts (D). Phase contrast images of senescence associated β-Galactosidase staining for top seven molecules that induced senescence and SLO and SSO combinations with neonatal fibroblasts (E), and quantification results for percentage of positive cells (all numbers across replicates pooled) and divided to highly expression and moderate expression classes based on intensity of staining (F). (*: p<0.05, **: p<0.01, ***: p<0.001 one-way ANOVA with Dunnett's multiple comparison test). Scale bar = 100um.

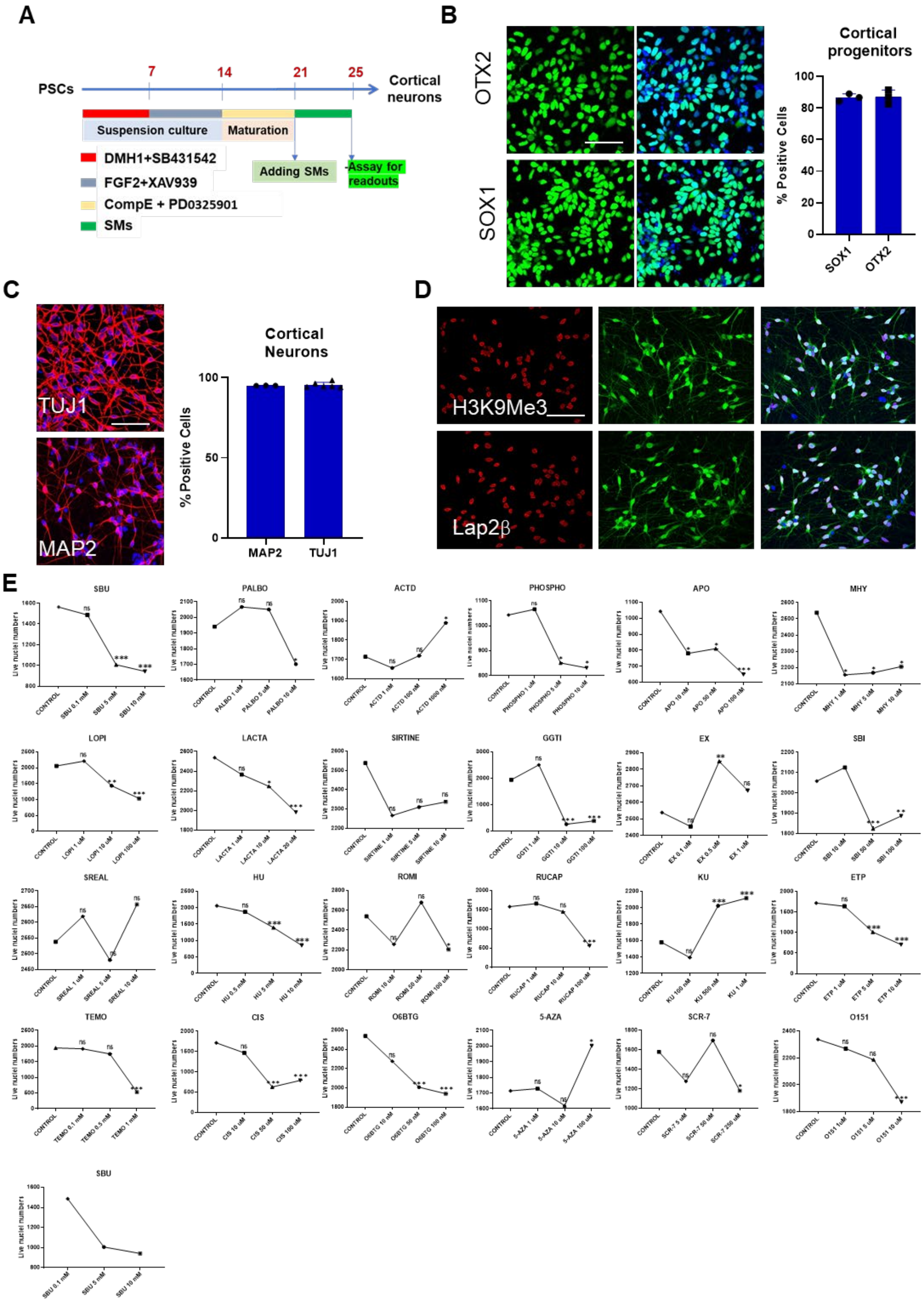


**Figure S2. related to figure 3- differentiation of cortical neurons from H9-GFP ESCs and characterization for expression of neuronal markers.**

Differentiation protocol used for generating cortical neurons from H9-GFP stem cells (A). Immunostaining images for SOX1 and OTX2 in day-14 cortical progenitors and quantification for proportion of positive cells Scale bar=50 μm (B). Representative immunostaining images for day-21 cortical neurons expressing TUJ1 (TUBB3) and MAP2 proteins in red and nucleus stained with Hoechst in blue (Scale bar=100 μm) and quantification for percentage of positive neurons (C). Immunostaining images for H3K9Me3 and Lap2β proteins in day-21 GFP labeled cortical neurons, Scale bar=100 μm (D). Cell toxicity assay for different doses of 25 small molecules with cortical neurons (E). (ns: not significant, *: p<0.05, **: p<0.01, ***: p<0.001 one-way ANOVA with Dunnett's multiple comparison test).

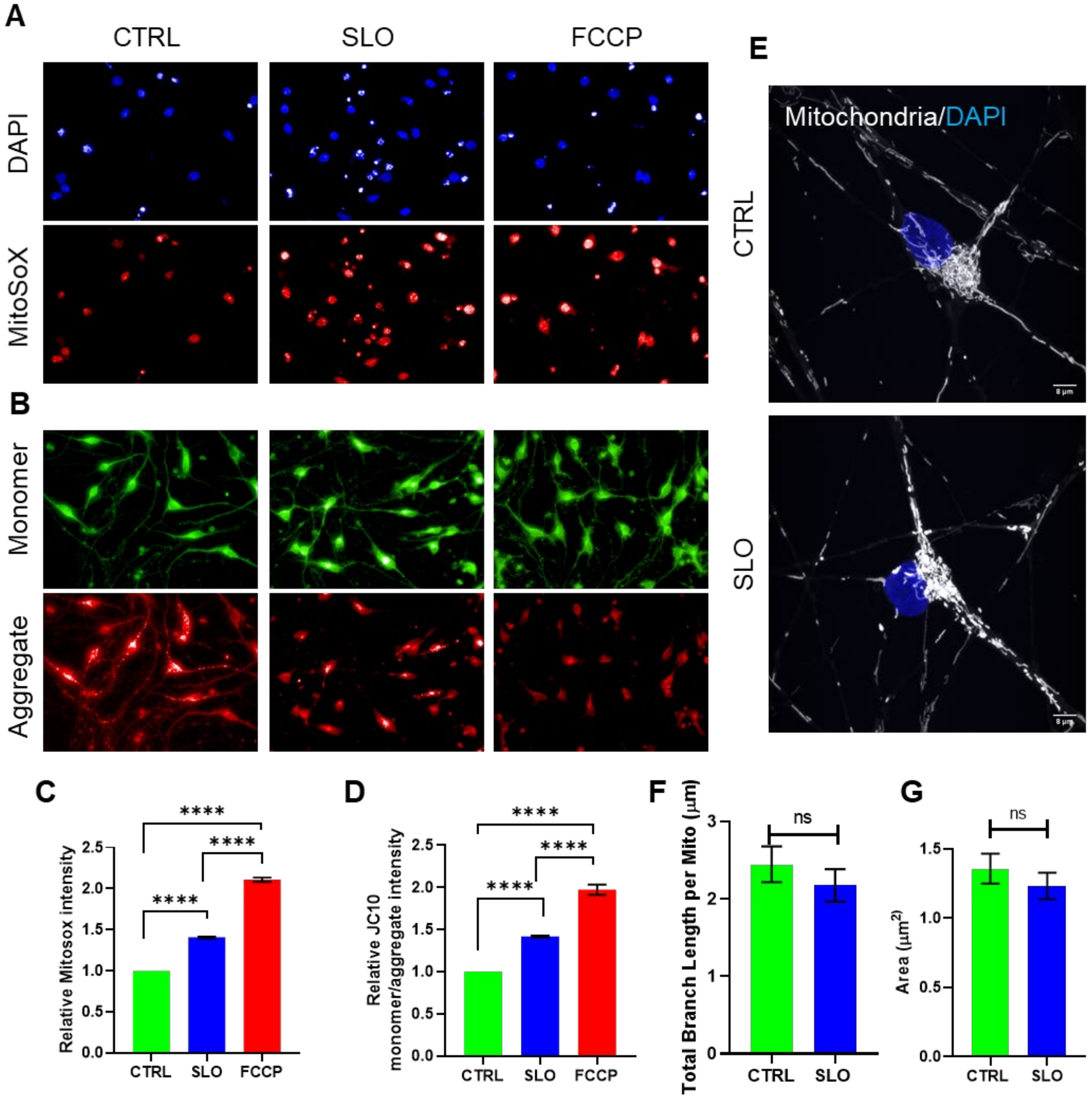

**Figure S3. MitoSoX and Mitochondrial membrane potential in SLO treated cortical neurons.** Representative images of cortical neurons at day 25 for MitoSoX staining (A) and JC-10 fluorescence (Top- Monomer (green), Bottom- Aggregate (red) from Control, SLO and FCCP treated cortical neurons Scale bar= 50µm (B). Statistical analysis of relative MitoSoX intensity from Control, SLO and FCCP treated neurons (C). Statistical analysis of relative JC10 Monomer to Aggregate intensity from Control, SLO and FCCP treated neurons (Low monomer to aggregate ratio means high mitochondrial membrane potential, high monomer to aggregate ratio means low mitochondrial membrane potential) (D). Represented images of Mitotracker stained mitochondria in cortical neurons (E) and quantification results for mitochondrial length (F) and mitochondrial area in SLO treated neurons versus control neurons (G). Data was quantified using 15,000 cells per group from two independent experiments. Statistical analysis was performed using One-way ANOVA, Tukey post-hoc test (****- $P<0.001$).

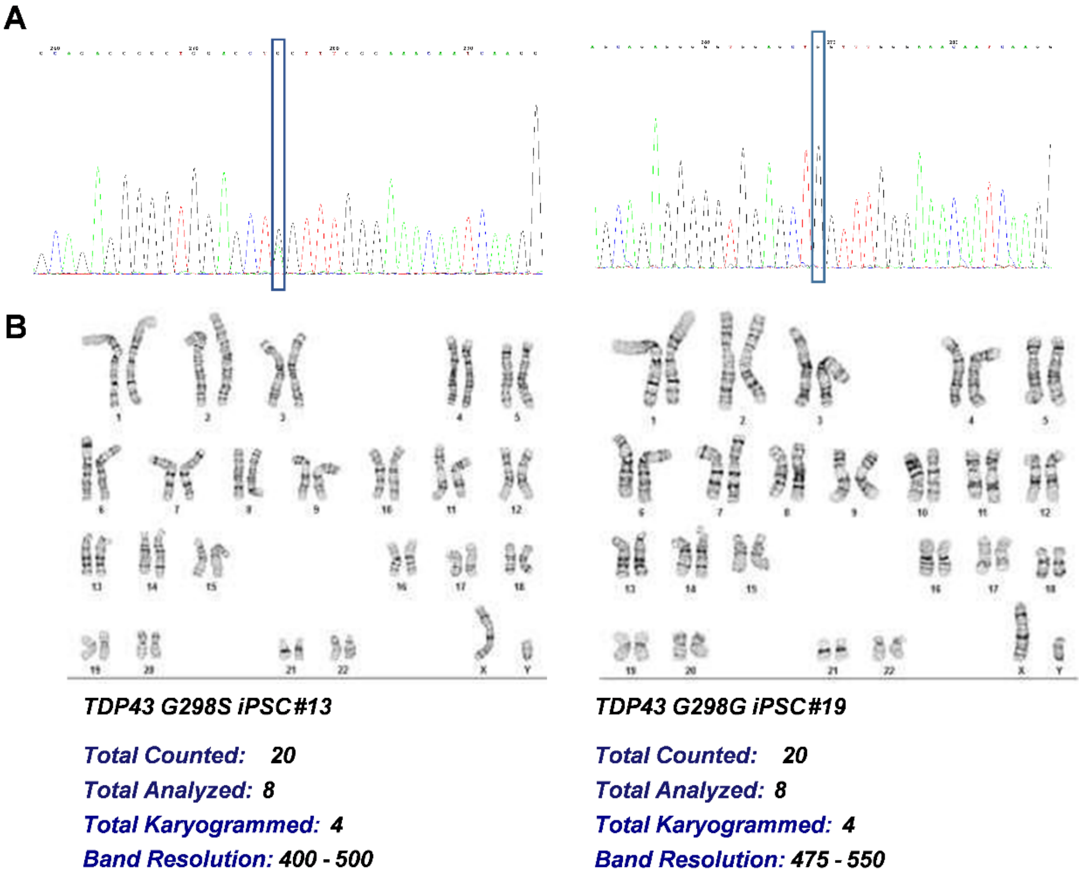


C

| STR Locus | STR Genotype Repeat # | STR Genotype |
|---|---|---|
| FGA | 16–18,18.2,19,19.2,20,20.2,21,21.2,22, 22.2, 23, 23.2, 24, 24.2, 25, 25.2, 26–30, 31.2, 43.2, 44.2,45.2, 46.2 | 19,21 |
| TPOX | 6-13 | 9,11 |
| D8S1179 | 7-18 | 10,12 |
| vWA | 10-22 | 17,17 |
| Amelogenin | X,Y | X,Y |
| Penta_D | 2.2, 3.2, 5, 7-17 | 11,12 |
| CSF1PO | 6-15 | 10,13 |
| D16S539 | 5, 8-15 | 9,11 |
| D7S820 | 6-14 | 8,12 |
| D13S317 | 7-15 | 11,12 |
| D5S818 | 7-16 | 11,14 |
| Penta_E | 5-24 | 7,16 |
| D18S51 | 8-10, 10.2, 11-13, 13.2, 14-27 | 12,16 |
| D21S11 | 24,24.2,25,25.2,26-28,28.2,29,29.2, 30, 30.2,31, 31.2,32,32.2,33,33.2, 34,34.2,35,35.2,36 -38 | 30,31 |
| TH01 | 4-9,9.3,10-11,13.3 | 6,9.3 |
| D3S1358 | 12-20 | 15,15 |

| STR Locus | STR Genotype Repeat # | STR Genotype |
|---|---|---|
| FGA | 16 18,18.2,19,19.2,20,20.2,21,21.2,22, 22.2, 23, 23.2, 24, 24.2, 25, 25.2, 26 30, 31.2, 43.2, 44.2,45.2, 46.2 | 19,21 |
| TPOX | 6-13 | 9,11 |
| D8S1179 | 7-18 | 10,12 |
| vWA | 10-22 | 17,17 |
| Amelogenin | X,Y | X,Y |
| Penta_D | 2.2, 3.2, 5, 7-17 | 11,12 |
| CSF1PO | 6-15 | 10,13 |
| D16S539 | 5, 8-15 | 9,11 |
| D7S820 | 6-14 | 8,12 |
| D13S317 | 7-15 | 11,12 |
| D5S818 | 7-16 | 11,14 |
| Penta_E | 5-24 | 7,16 |
| D18S51 | 8-10, 10.2, 11-13, 13.2, 14-27 | 12,16 |
| D21S11 | 24,24.2,25,25.2,26-28,28.2,29,29.2, 30, 30.2,31, 31.2,32,32.2,33,33.2, 34,34.2,35,35.2,36 -38 | 30,31 |
| TH01 | 4-9,9.3,10-11,13.3 | 6,9.3 |
| D3S1358 | 12-20 | 15,15 |

**Figure S4. Related to figure 6- Characterization of TDP43 G298S and TDP43 G298G iPSCs.**

Sanger sequencing result for both mutant (left panel) and corrected (right panel) (isogenic control) cell lines. Karyotype analysis for mutant (left) and corrected (right) cell lines (B). STR analysis for both cell lines were done for selected loci depicted in C.

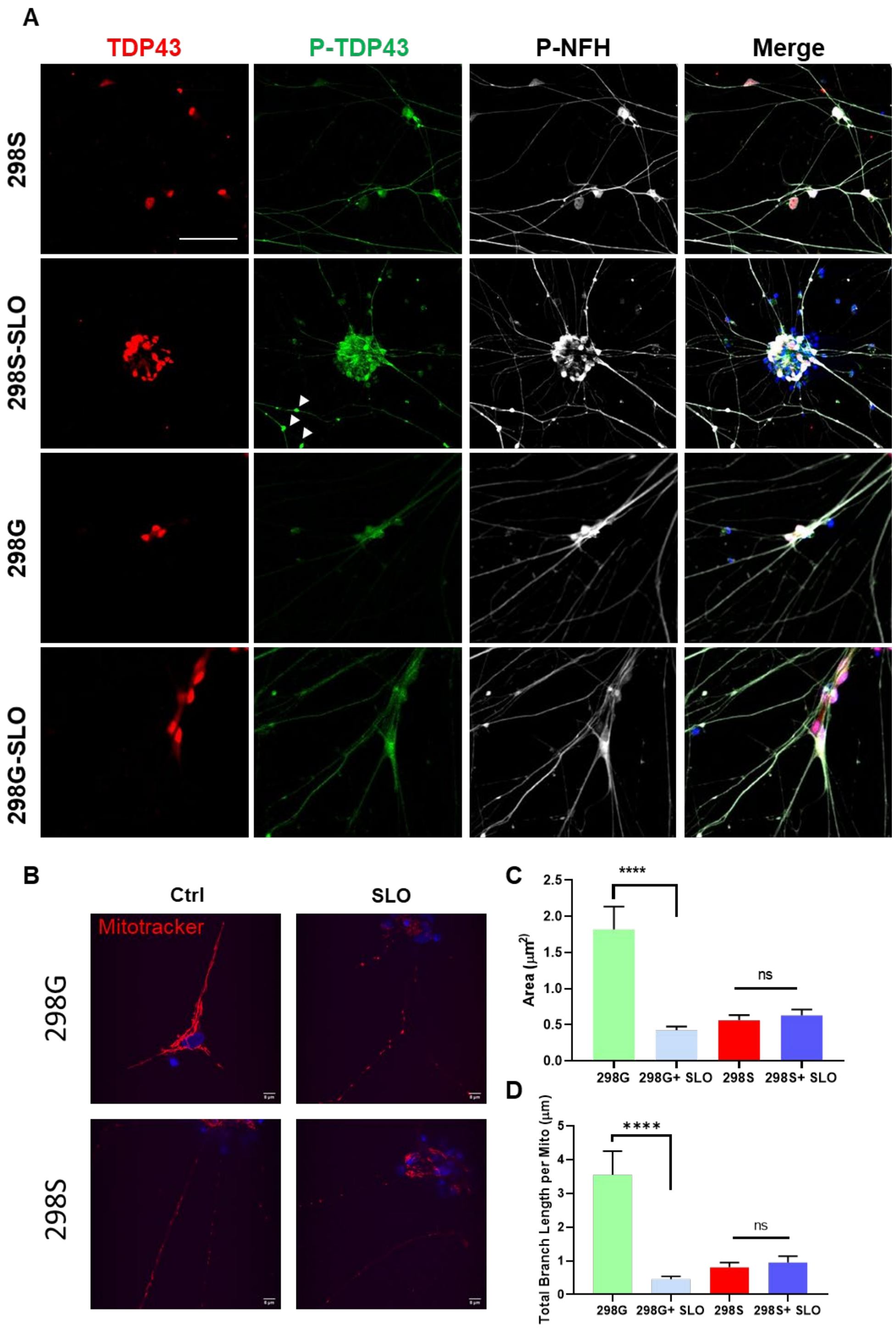


**Figure S5. Related to Figure 6- ALS MNs treated with SLO molecules shows signs of neurodegeneration and protein phosphorylation.** Immunostaining images of phosphor-TDP43 protein following SLO treatment in 298S mutant cells and 298G healthy control cells, arrow heads are pointed at p-TDP43 positive neurite swellings in mutant cells (A, for quantification see the Figure 6H). (Scale bar=100 μm). MNs stained using Mitotracker red to visualize morphological changes in SLO treated cells (B) (Scale bar=8 μm), and graphs of quantified data for mitochondrial area (C) and total branch length for each mitochondria (D). ( *: $p<0.05$, ***: $p<0.001$ one-way ANOVA with Dunnett's multiple comparison test).

**Table S1: List of the small molecules and their concentration for final screen in neurons.**

| Name | Function | Working Concentration | Name | Function | Working Concentration |
|---|---|---|---|---|---|
| **WZ4003** | AMPK inhibitor | 2μM | **SirReal2** | Sirt2 inhibitor | 1μM |
| **MHY1485** | mTOR activator | 2μM | **Rucaparib** | PARP1 inhibitor | 1μM |
| **Fumonisin B1** | AKT activators | 5μM | **Temozolomide** | DNA alkylation | 100μM |
| **Sirtinol** | Sirtuin inhibitors | 5μM | **Lactacystin** | irreversible proteasome inhibitor | 4nM |
| **SBI-0206965** | Autophagy inhibitor | 10μM | **KU-60019** | ATM inhibitor | 100nM |
| **Romidepsin** | HDAC1,2 inhibitor | 10pM | **5-AZA-20-DEOXYCYTIDINE** | DNA methyltransferase inhibitor | 1μM |
| **Etoposid** | Topo II inhibitor | 2μM | **Actinomycin D** | inhibiting DNA-primed RNA synthesis | 10nM |
| **Lomeguatrib-O6BTG** | MGMT inhibitor | 10nM | **Cisplatin** | Topo I,II inhibitor | 10μM |
| **O151** | DNA Glycosylase-1 inhibitor | 1μM | **SCR-7** | Ligase V inhibitor | 5μM |
| **Palbociclib** | CDK4/6 inhibitor | 1μM | **Phosphoramidon** | metalloendopeptidase inhibitor | 1μM |
| **Apo866** | NAD biosynthesis inhibitor | 10nM | **Lopinavir** | HIV protease inhibitor | 1μM |
| **Hydroxyurea** | DNA synthesis stress inducer | 500μM | **GGTI298** | geranylgeranyltransferase I (GGTase I) inhibitor. | 1μM |
| **EX-527** | Sirt1 inhibitor | 100nM | **Sodium Butyrate** | histone deacetylase inhibitor I, II | 100μM |
| **SMER28** | Autophagy activator | 1μM | **Edaravone** | Radical scavenger | 1μM |
| **Tat-Beclin** | Autophagy activator | 100nM | **Amiodarone** | $K^+$ channel blocker | 5μM |
| **STF-62247** | Autophagy activator | 1μM | **Flubendazole** | Autophagy activator | 1μM |
